# Erasing motion: Scrambling direction selectivity in visual cortex during saccades

**DOI:** 10.1101/2021.03.30.437338

**Authors:** Satoru K. Miura, Massimo Scanziani

## Abstract

Distinguishing sensory stimuli caused by changes in the environment from those caused by an animal’s own actions is a hallmark of sensory processing. Saccades are rapid eye movements that shift the image on the retina. How visual systems differentiate motion of the image induced by saccades from actual motion in the environment is not fully understood. Here, we discovered that in mouse primary visual cortex (V1), the two types of motion evoke distinct activity patterns. This is because during saccades, V1 combines the visual input with a strong non-visual input arriving from the thalamic pulvinar nucleus. Thus, the pulvinar input ensures differential V1 responses to external and self-generated motion and may allow the animal to interpret sensory stimuli within the context of its actions.

## Main Text

Sensory stimuli are often generated by the animal’s own movements, and nervous systems have evolved mechanisms to distinguish those self-generated stimuli from externally generated ones (*1*). Prime examples are saccades, rapid eye movements that induce a fast displacement of the visual scene on the retina. They are common in animals across phyla, including animals without fovea like rodents, and they contribute to shifts of the gaze (*2–6*). Behavioral studies indicate that such saccade-induced motion of the visual scene is distinguished by the subjects from motion occurring in the environment (*7–12*).

How visual systems distinguish the two types of motion, despite similar shifts of the image on the retina, has been a long-standing question. A non-visual, extra-retinal signal occurring around the time of saccades has been proposed to play a key role. This non-visual signal is believed to be transmitted to specific nodes along the visual pathway, where it interacts with the neural responses to saccade-induced motion of the visual scene. One model proposes that the non-visual signal alters the pattern of the responses to motion (*13*), such that the representation of saccade induced motion of the visual scene is distinct from that of actual motion in the environment. An alternate model proposes that it suppresses the responses to the saccade-induced motion (*14–17*). These models are not mutually exclusive. However, the origin of the non-visual signal to the visual cortex, what it encodes, and how it impacts the neural representation of the motion induced by saccades is not known.

### Saccade direction preference in V1

We recorded the response of V1 neurons to saccades in unrestrained, freely moving mice with chronically implanted extracellular electrodes while tracking the movement of the eye contralateral to the recorded hemisphere with a head-mounted miniature camera (Fig. 1A - C). Saccades occurred in all directions yet were biased along the horizontal over vertical axis (13708 horizontal and 6913 vertical saccades from 5 animals; binomial test, *p* < 0.0001; Fig. 1D). They had a frequency of 45.3 ± 6.4 events/min, mean amplitude of 18.8 ± 1.0°, and mean 10 - 90% rise time of 25.9 ± 0.9 ms, resulting in average speed of 709 ± 31° /s [average ± standard deviation (s.d.) of 5 mice]. A large fraction of V1 neurons showed time-locked responses to saccades (105 out of 169, 5 mice; Fig. 1E, fig. S1; see Methods for definition). Strikingly, half of those neurons discriminated saccade direction (53 out of 105; Fig. 1E – H, fig. S1; discriminability was calculated based on receiver-operating characteristic analysis; see Methods), and their responses both preceded and outlasted saccades by several tens of milliseconds, as shown by the peri-event time histogram (PETH; Fig 1H). Their direction selectivity index averaged 0.54 ± 0.24 (average ± s.d.; fig S1; see Methods), and their preferred directions were unevenly distributed (Rao’s spacing test, *p* < 0.001; Fig. 1G). Thus, neurons in V1 respond to saccades in a direction selective manner.

**Fig. 1.**
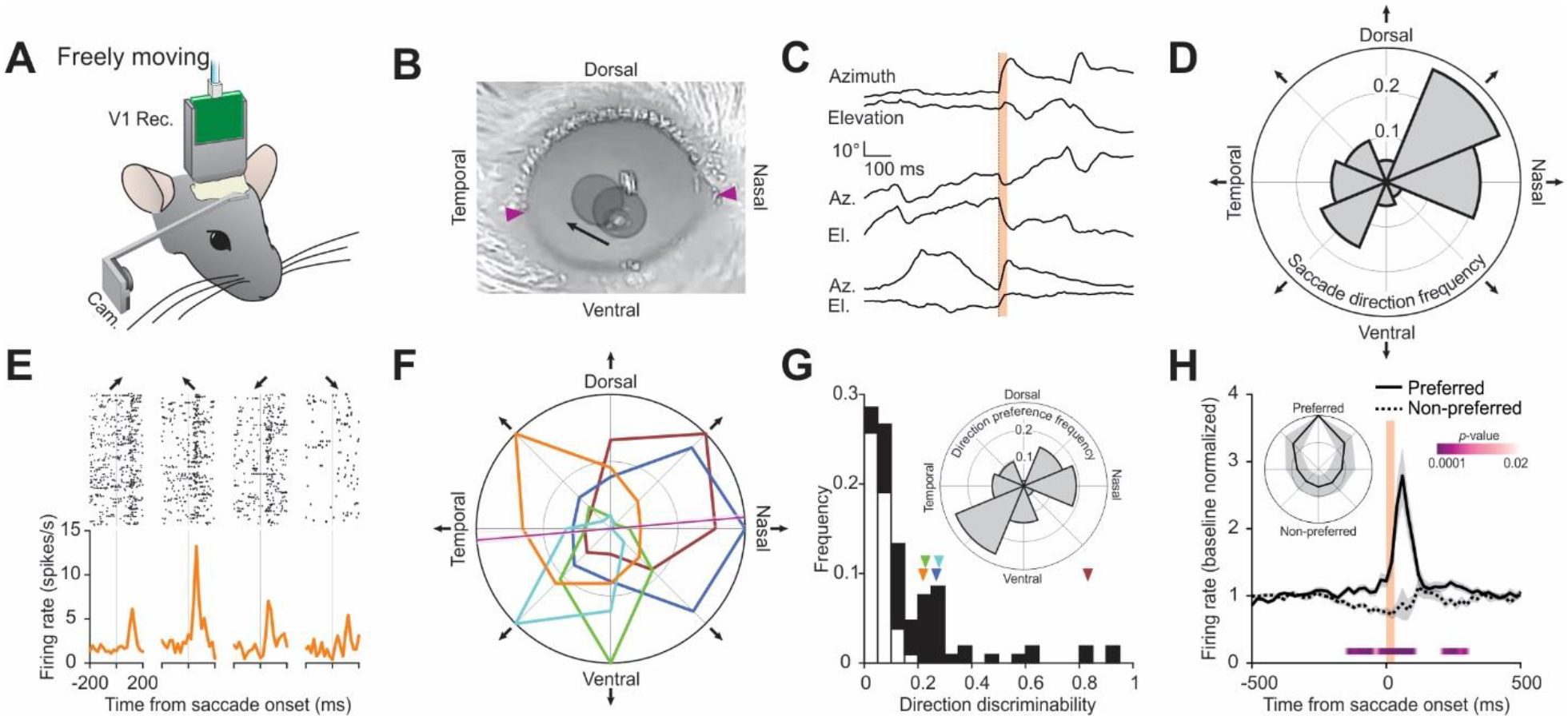
V1 neurons are tuned to saccade direction. (**A**) Schematic of experimental setup in freely moving mice. (**B**) Overlay of two snapshots of the right eye, taken before and after a dorso-temporal saccade. Arrow indicates direction of saccade. Pupils are overlaid with gray circles. Magenta arrowheads, lateral (left) and medial (right) commissures. (**C**) Traces of the eye position for three example saccades. Vertical orange bar, 0 - 90% rise time of saccades (31 ms). In each pair: top trace, azimuth (up is nasal); bottom trace, elevation (up is dorsal). (**D**) Polar histogram showing saccade direction frequency. Average of five animals, normalized. (**E**) Example V1 neuron showing saccade direction preference for dorso-temporal direction. Raster plots (top) and peri-event time histogram (PETH; bottom). The arrows on the top indicate the direction of saccades as shown in (F). (**F**) Polar plot of five example saccade direction selective neurons, normalized to their maximum response. Orange, example neuron in (E). Magenta lines indicate the angle of the axis connecting the lateral and medial commissures [arrowheads in (B); lighter shade, individual animal; darker shade, average]. (**G**) Direction discriminability of saccade responsive neurons (calculated based on receiver operating characteristic of spike frequency distribution, 0 indicates no discriminability, 1 indicates perfect discriminability; see Methods). Black, direction selective neurons (*n* = 53 neurons); white, non-selective (*n* = 52 neurons, 5 mice). Arrowheads show discriminability of the example neurons in (F). Inset, polar histogram of preferred direction frequency. (**H**) Average PETH of saccade direction selective neurons (*n* = 53 neurons, 5 mice) for preferred and non-preferred (i.e. opposite) directions. Baseline normalized. Shaded area, average ± standard error of the mean (s.e.m.). Vertical orange bar, 0 - 90% rise time of saccades (31 ms). *P*-values, comparison of activity for preferred and non-preferred directions in 20-ms bins (Wilcoxon signed-rank test, one-tailed, only significant values shown; see Methods). Inset, polar plot of average response of direction selective neurons, aligned for the preferred direction. On average, direction selective neurons fired 3.0 times more in response to saccades in the preferred than the non-preferred direction. Shaded area, standard deviation (s.d.).

### Scrambling of stimulus direction preference by saccades

The response to saccades of V1 neurons as well as their saccade direction preference may simply result from the saccade induced motion of the image on the retina. Indeed, V1 neurons respond robustly to moving visual stimuli and often do so in a direction selective manner (*18–20*).

To assess whether neuronal activity in response to saccades is the direct consequence of the motion of the visual scene on the retina, we performed extracellular recordings in V1 of head-fixed, awake mice. The head-fixed configuration allowed us to more precisely control the visual environment of the animal. We presented a stationary vertical sinusoidal grating on a monitor placed in the visual field contralateral to the recorded hemisphere and tracked the movements of the corresponding eye (Fig. 2A, B). We recorded the activity in response to shifts of the grating on the retina resulting from spontaneous saccades and compared it to the activity in response to pseudo-saccades. Pseudo-saccades are shifts of the grating on the monitor in the absence of the animal’s eye movements and are designed to approximate the shifts resulting from real saccades. Saccades in head-fixed animals occurred almost exclusively along the horizontal axis in either the nasal or temporal direction, consistent with previous reports (*21, 22*) (Fig. 2C, D), had a frequency of 3.3 ± 0.3 saccades/min (nasal, 2.0 ± 0.2; temporal 1.3 ± 0.2), mean amplitude of 10.9 ± 1.9° (nasal, 13.1 ± 2.8; temporal, 7.5 ± 1.1; fig. S2), and mean 10 - 90% risetime of 23.0 ± 1.5 ms (nasal, 19.4 ± 1.7; temporal, 28.3 ± 3.1), resulting in average speed of 472 ± 100° /s [nasal, 601 ± 158; temporal, 274 ± 69; all stats average ± standard deviation (s.d.) of 4 mice].

**Fig. 2.**
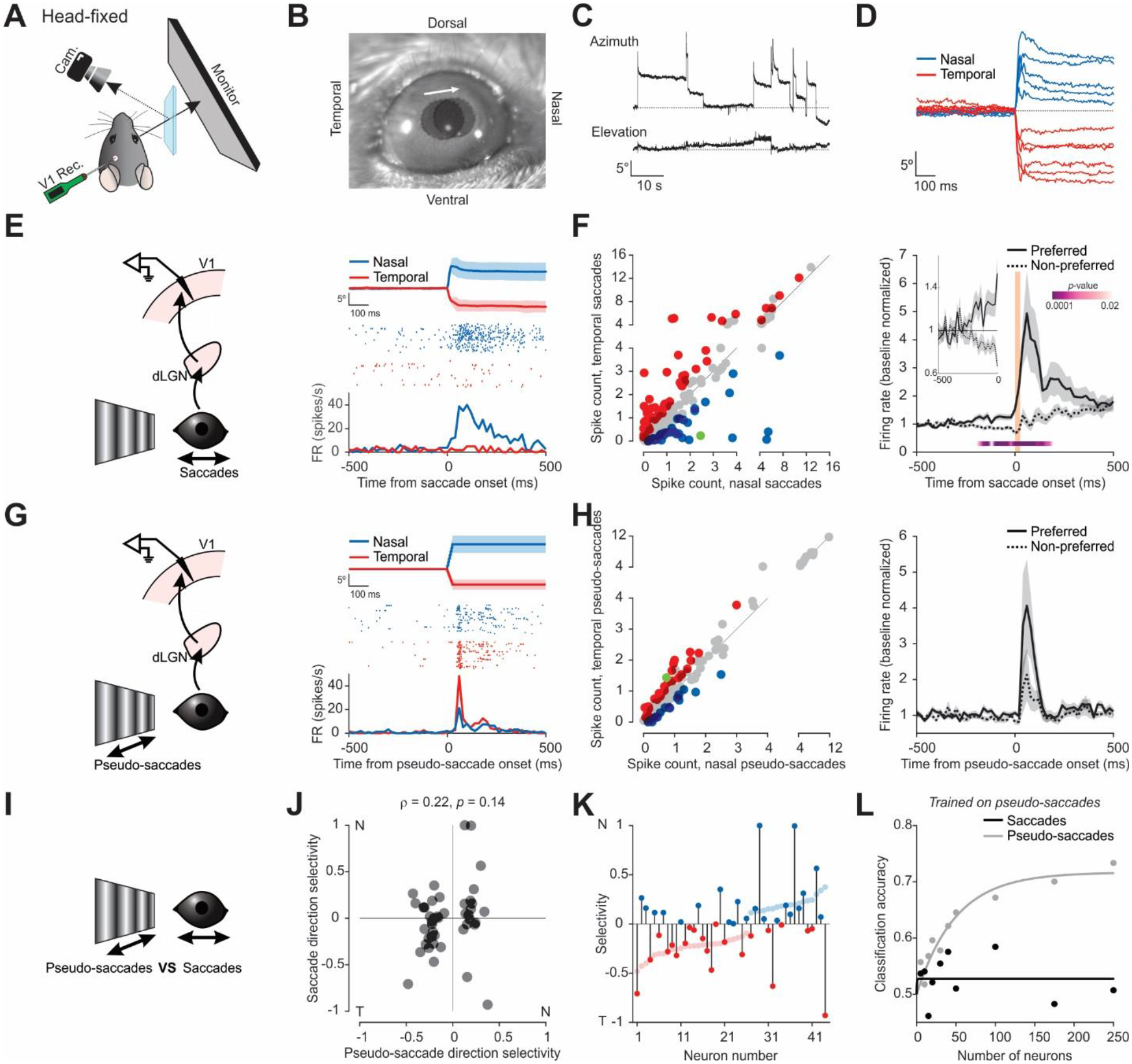
Scrambling of stimulus direction selectivity in V1 during saccades. (**A**) Schematic of experimental setup in head-fixed mice. (**B**) Overlay of two snapshots of the right eye, taken before and after a nasal saccade. Pupils are overlaid with gray circles. Arrow indicates direction of saccade. (**C**) Example traces of the eye position over 1-minute. Top trace, azimuth (up is nasal); bottom trace, elevation (up is dorsal). (**D**) Example traces of relative azimuthal eye position for nasal and temporal saccades. (**E**) Left, schematic of V1 recording during saccades on a vertical grating. Right, example neuron preferring nasal saccades. Average eye position for nasal and temporal saccades (top; shaded area, average ± standard deviation), raster plots (center), and PETH (bottom). (**F**) Left, scatter plot of the response to nasal and temporal saccades (average spike count in a 100 ms window from saccade onset), for all responsive neurons (*n* = 189 neurons, 4 mice; see Methods). Blue, prefer nasal saccades; red, prefer temporal saccades; gray, no statistical difference; green, example neuron in (E). Significance determined by rank-sum test, critical value 0.05. Right, average PETH of discriminating neurons (colored data points in left scatter plot; *n* = 97 neurons, 4 mice), for preferred and non-preferred saccade directions. Baseline normalized. Shaded area, average ± s.e.m. Vertical orange bar, 0 - 90% rise time of saccades for reference (26 ms). *P*-values, comparison of activity for preferred and non-preferred saccade directions in 20-ms bins (Wilcoxon signed-rank test, one-tailed). Inset, magnified, note separation prior to saccade onset. (**G**) Left, schematic of V1 recording during pseudo-saccades. Right, example neuron preferring temporal pseudo-saccades (otherwise as in (E)). (**H**) Left, scatter plot of response to nasal and temporal pseudo-saccades (*n* = 157 neurons, 4 mice). Right, average PETH of discriminating neurons (*n* = 44 neurons, 4 mice; otherwise as in (F)). (**I**) Comparison of the response to saccades and pseudo-saccades. (**J**) Scatter plot of direction selectivity for real and pseudo-saccades for all neurons discriminating pseudo-saccade directions. Negative values, prefers temporal; positive values, prefers nasal (see Methods). Note poor correlation. Pearson ρ = 0.22, *p* = 0.14, *n* = 44 neurons, 4 mice. (**K**) Lighter shade, neurons sorted according to their direction selectivity for pseudo-saccades. Darker shade, direction selectivity of the same neurons for real saccades. Note the scrambling, i.e., several neurons change direction preference. (**L**) Classification accuracy of direction of motion (nasal or temporal) of a linear classifier trained on pseudo-saccades and tested on pseudo-saccades and real saccades, as a function of the number of neurons included in the analysis (see Methods). Lines, exponential fits. Note reduced accuracy on real saccades.

Similar to freely moving animals, the activity of the majority of V1 neurons (∼60%) sampled across all layers was significantly modulated within the first 100 ms from saccade onset (189 out of 317 neurons, 4 mice; see Methods for definition; fig. S3), and more than half of the responding neurons exhibited a significant direction preference for either nasal or temporal saccades (97 out of 189; Fig. 2E, F). The direction preference was observed in regular-spiking (putative excitatory) as well as fast-spiking (putative inhibitory) neurons (fig. S3). Furthermore, again similar to freely moving animals, the activity and the directional preference both preceded and outlasted the saccade duration by several tens of milliseconds (Fig. 2F, right). Thus, also in head-fixed animals, saccades elicit strong, direction selective responses in V1. To compare the responses between real and pseudo-saccades, we presented, to the same mice in the same session, pseudo-saccades with a rise-time of 25 ms and whose amplitudes (i.e., horizontal shift of the grating) ranged from 3.0° to 24.6° (see Methods). Shifts of the grating in the nasal direction were termed temporal pseudo-saccades because they generated a shift of the image on the retina in the same direction as that generated by real temporal saccades. Conversely, temporal shifts of the grating were termed nasal pseudo-saccades. For the analysis, we selected pseudo-saccades whose amplitudes and directions were matched to those of the real saccades performed by each animal during the recording session [average amplitude 10.9 ± 1.8° (nasal, 13.1 ± 2.7; temporal, 7.6 ± 1.0; fig. S2), average speed 437 ± 73° /s (nasal, 522 ± 108; temporal, 303 ± 39; all stats average ± s.d. of 4 mice); see Methods]. A large fraction of V1 neurons (50%, 157 out of 317) responded to pseudo-saccades, and 28% of these neurons (44 out of 157) showed a preference for the nasal or temporal direction (Fig. 2G, H; fig. S3).

If saccade-induced responses in V1 result from the shift of the grating on the retina, the direction preference of neurons to saccades and pseudo-saccades should correlate. In marked contrast to this expectation, there was a lack of correlation between the direction preference to real and pseudo-saccades of V1 neurons [Pearson correlation coefficient (ρ) = 0.22, *p* = 0.14, *n* = 44; Fig. 2J, K]. Among the neurons that discriminated the direction of pseudo-saccades, 41% reversed direction preference in response to real saccades (18 out of 44), and, for the rest of the neurons, direction selectivity was decreased or enhanced, irrespective of the pseudo-saccade direction preference and selectivity (Fig. 2J, K). Furthermore, among the neurons that responded but did not discriminate the direction of pseudo-saccades, 35% became direction selective in response to real saccades (40 out of 113; fig. S4). Thus, saccades and pseudo-saccades induce distinct patterns of activity in V1, suggesting that the response of V1 neurons to saccades does not result solely from the shift of the visual scene on the retina.

Despite the poor correlation, the distributions of direction preference for real and pseudo-saccades were similar, as if direction preference had been randomly reassigned using the original probability distribution (22 out of 44 prefer nasal saccades, 18 out of 44 prefer nasal pseudo-saccades; Z-test, *p* = 0.39). We therefore refer to the impact of saccades on direction preference as “scrambling”.

Consistent with the poor correlation between the response to real and pseudo-saccades, a linear classifier of motion direction (i.e., nasal or temporal) trained on V1 population response to pseudo-saccades was unable to generalize to the response to real saccades. The accuracy of the classifier (trained over a time window of 100 ms from pseudo-saccade onset), which decodes the direction of pseudo-saccades well above chance, fell to chance for real saccades (73.3 ± 5.0% accuracy ± s.d. for pseudo-saccades with 250 neurons, chance level 48.5 ± 6.8%, Wilcoxon rank-sum test, two-tail, *p* < 0.0001, *n* = 300; 50.7 ± 7.1% for real saccades, chance level 50.0 ± 7.1%, Wilcoxon rank-sum test, two-tail, *p* = 0.32, *n* = 300; see Methods; Fig. 2L). These results show that saccades scramble direction selectivity in V1.

### Non-visual, saccade mediated responses in V1

Why is the response of V1 neurons to real saccades so different from the response to pseudo-saccades? We hypothesized that, around the time of saccades, a non-visual signal enters V1 thereby altering the response to the visual signal arriving from the retina. Consistent with this hypothesis, in neurons showing saccade direction preference, the PETHs for preferred and non-preferred saccade directions diverged ∼150 ms before saccade onset, both in freely moving and head-fixed animals (140 ms for freely moving, 180 ms for head-fixed; Fig. 1H, Fig. 2F, right). Given that the modulation of V1 activity occurs before the saccadic eye movement, it cannot be accounted for, at least in head-fixed animals, by a change in the visual scene on the retina, suggesting the presence of a non-visual signal.

To isolate this putative non-visual signal, we used two different approaches, both aimed at preventing changes in retinal activity during saccades. We recorded the response of V1 neurons to saccades made either in front of a gray screen covering a large portion of the visual field (i.e. a visual scene where the luminance is homogeneous in space, see Methods; Fig. 3A, B) or with tetrodotoxin (TTX) injected in both eyes to block retinal activity (Fig. 3C, D; distribution of saccade amplitudes was little affected by the TTX injection compared to control, fig. S2). In both cases, saccades still triggered strong, directionally selective responses in V1. With the gray screen, 55% of all neurons responded to saccades, of which 63% discriminated the direction, thus resembling saccades made on the vertical grating (174 responsive, 109 discriminating out of 317 total, 4 mice; Fig. 3A, B). Furthermore, the PETH for the preferred and non-preferred directions diverged prior to saccade onset [200 ms window before onset, 0.2 ± 0.1 Hz evoked firing rate (FR) ± s.e.m. for preferred, −0.1 ± 0.03 Hz for non-preferred; Wilcoxon signed-rank test, one-tailed, *p* < 0.0001, *n* = 109; Fig. 3B, right]. Similarly, in TTX-blinded animals, about half of the neurons in V1 responded to saccades, of which 64% discriminated the direction (100 responsive, 64 discriminating out of 203 total, 8 mice; Fig. 3C, D; fig. S5), and the PETH for the preferred and non-preferred directions diverged well before saccade onset (200 ms window before onset, 1.0 ± 0.3 Hz evoked FR ± s.e.m. for preferred, −0.4 ± 0.2 Hz for non-preferred; Wilcoxon signed-rank test, one-tailed, *p* = 3.3 x 10^-4^, *n* = 64; Fig. 3D, right). The impact of saccades on neuronal activity was layer-dependent, showing a gradient of increasing excitability and discriminability as a function of depth (fig. S5). The block of visual input in TTX-blinded animals was verified by the complete absence of responses in V1 to visual stimuli (Fig. 3C; fig. S6). This complete block allowed us to estimate the site of entry of the non-visual input by performing a current source density analysis of the saccade-triggered local field potential (LFP). The analysis revealed a strong sink in the supragranular layers of V1, distinct from the initial sink in layer 4 in response to a visual input (fig. S5). These data demonstrate that mouse V1 receives a non-visual input that targets the superficial layers and that carries saccade direction information.

**Fig. 3.**
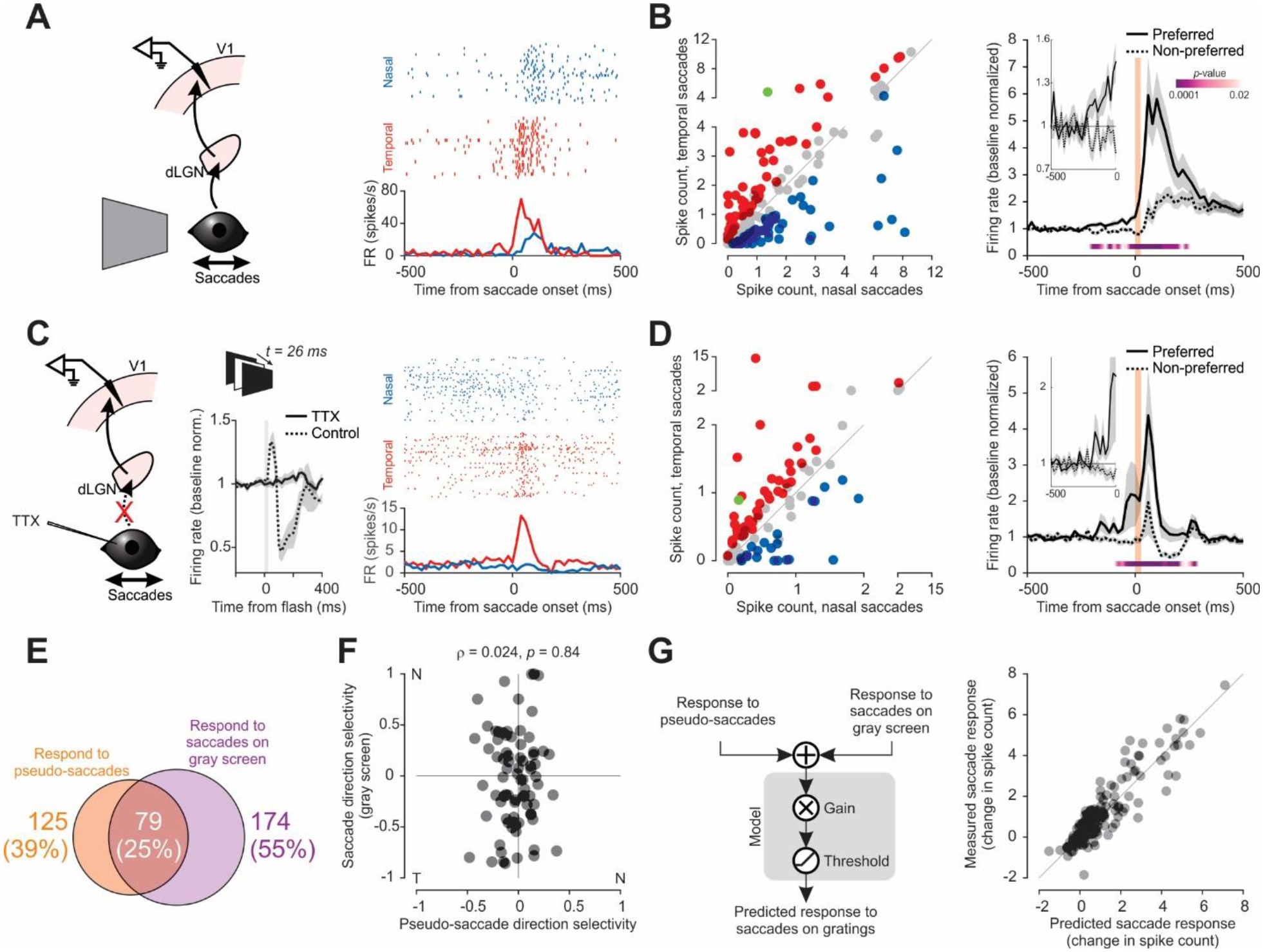
V1 receives non-visual direction selective input during saccades. (**A**) Left, schematic of V1 recording during saccades on a gray screen. Right, example neuron preferring temporal saccades. Raster plots (top) and PETH (bottom). (**B**) Left, scatter plot of the response to nasal and temporal saccades for all responsive neurons. Blue, prefers nasal saccades; red, prefers temporal saccades; gray, no statistical difference; green, example neuron in (A). Right, Average PETH of discriminating neurons (colored data points in left scatter plot) for preferred and non-preferred saccade directions (*n* = 109 neurons, 4 mice). Inset, magnified, prior to saccade onset. Baseline normalized. Shaded area, average ± s.e.m. Vertical orange bar, 0 - 90% rise time of saccades (25 ms). *P*-values, comparison of activity for preferred and non-preferred saccade directions in 20-ms bins (Wilcoxon signed-rank test, one-tailed; only critical values shown). Inset, magnified, note separation prior to saccade onset. (**C**) Left, schematic of V1 recording during saccades in TTX-blinded animals. Center, average multi-unit responses to a brief full-field flash. Vertical gray bar indicates the flash duration (26 ms). Note the lack of response in TTX-blinded animals. *n* = 4 (control) and 8 (TTX-blinded) mice. See also fig. S6. Right, example neuron in a TTX-blinded animal, preferring temporal saccades. Raster plots (top) and PETH (bottom). (**D**) Same as in (B). Right, *n* = 64, 8 mice. 0 - 90% rise time of saccades, 27 ms. (**E**) Venn diagram of the number of neurons that respond to pseudo-saccades, saccades on a gray screen, and both. Percentages are out of the entire population. (**F**) Scatter plot of direction selectivity for pseudo-saccades (x-axis) against saccades on a gray screen (y-axis), for neurons that respond to both (79 neurons in (E)). Pearson ρ = 0.024, *p* = 0.84, *n* = 79 neurons, 4 mice. (**G**) Left, schematic of the linear regression-based model used to predict the number of spikes evoked by saccades on a vertical grating based on the response of neurons to pseudo-saccades and to saccades on a gray screen (see Methods). Right, predicted number of spikes (x-axis) plotted against the observed values (y-axis).

### Combining stimulus direction preference with saccade direction preference

To determine how the non-visual and the visual inputs interact in V1 during a saccade, we made a simple model based on linear regression. We used the responses to saccades in front of the gray screen as a proxy for the non-visual input and the responses to pseudo-saccades as a proxy for the visual input to predict the response to saccades made in front of a vertical grating. We based this analysis on 79 neurons that responded to both saccades on a gray screen and to pseudo-saccades (4 mice; Fig. 3E). Consistent with the above results (Fig. 2J, K), there was no correlation between the direction selectivity of these neurons to saccades on a gray screen and pseudo-saccades, indicating that the direction selectivity imparted by the visual and non-visual inputs to V1 neurons are independent (Pearson ρ = 0.024, *p* = 0.84, *n* = 79; Fig. 3F). Using a simple summation of the visual and non-visual inputs, this model explained 83% of the variance of the saccade response in front of a vertical grating (Fig. 3G; see Methods). The estimated gain on the combined input was 0.61 (*p* < 0.0001), suggesting a linear integration with reduced gain. In contrast, when using only the response from visual or from the non-visual inputs, the model explained only 32% or 69% of the variance, respectively (fig. S7). Taken together, these data suggest that the scrambling of the direction selectivity during a saccade results from the integration of visual and non-visual inputs whose direction selectivity are uncorrelated. To test this hypothesis, we proceeded to identify the source of the non-visual input. Silencing this source should eliminate the non-visual signal in V1 and prevent the scrambling of direction selectivity during saccades.

### The pulvinar nucleus is the source of saccade responses in V1

What is the source of the non-visual saccadic input to V1? We recorded from the dorsolateral geniculate nucleus of the thalamus (dLGN), the main source of afferent visual information to V1, to determine whether it is also the source of the non-visual input, since neurons in this structure have been shown previously to respond to saccades (*23–26*). In TTX-blinded animals, dLGN neurons responded to saccades, and their responses were selective for saccade direction (83 responsive, 49 discriminating, 174 total, 4 mice; Fig. 4A; fig. S8). To determine whether the dLGN is the source of the non-visual input to V1, we silenced the dLGN with muscimol injection in otherwise un-manipulated (i.e., non-blinded) animals. In contrast to the lack of visual responses in V1 confirming efficient silencing of the dLGN (fig. S9), V1 neurons still robustly responded to saccades and discriminated the two directions (106 responsive, 56 discriminating, total 139, 4 mice; Fig. 4B; fig. S8). These data show that the dLGN is not the main source of the non-visual saccade input to V1.

**Fig. 4.**
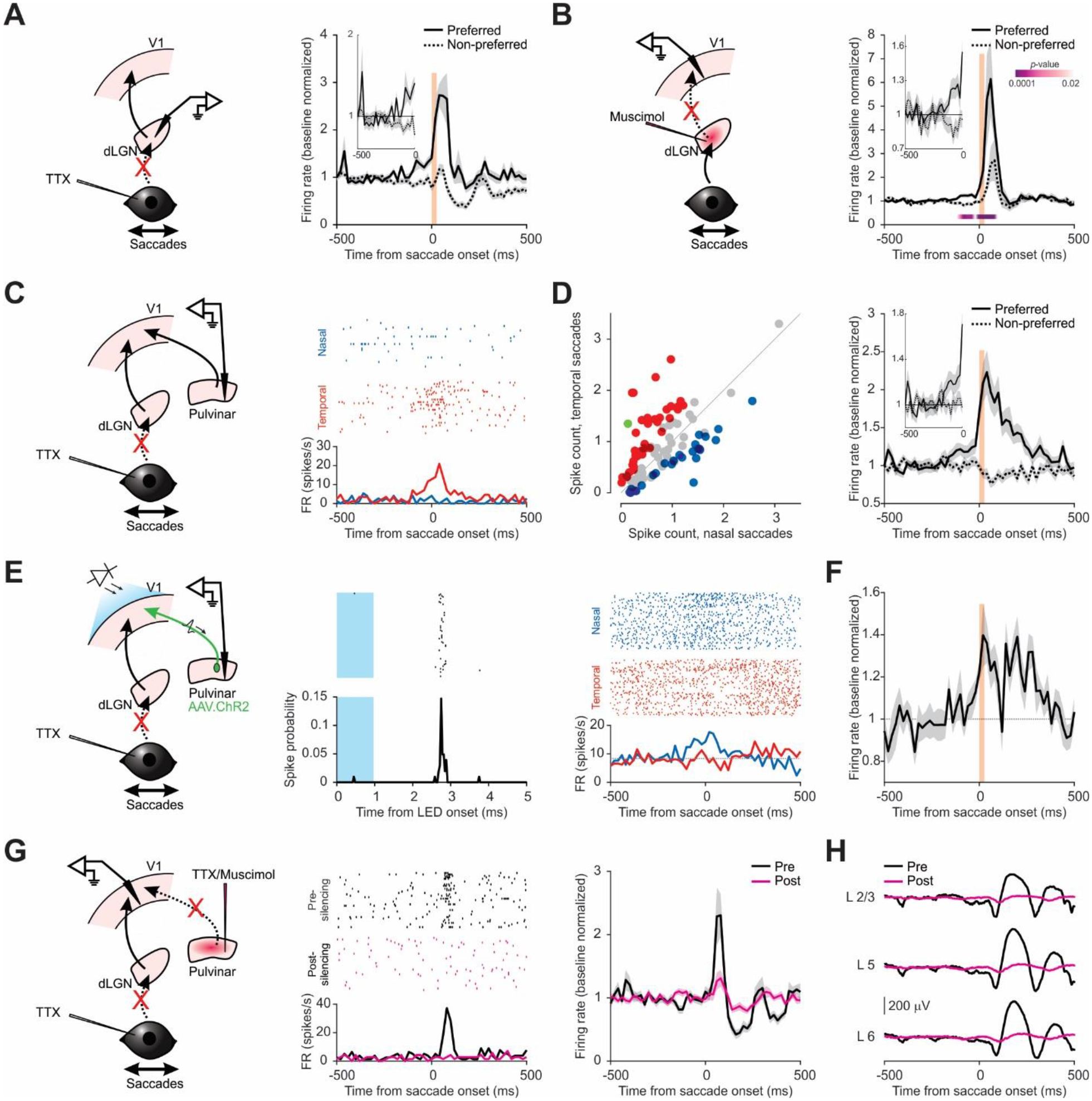
The pulvinar provides non-visual direction selective saccade input to V1. (**A**) Left, schematic of dLGN recording during saccades in TTX-blinded animals. Right, average PETH for preferred and non-preferred saccade directions (*n* = 49 neurons, 4 mice). Shaded area, average ± s.e.m. Vertical orange bar, 0 - 90% rise time of saccades (26 ms). Inset, magnified. (**B**) Left, schematic of V1 recording during saccades under dLGN silencing. Right, average PETH for preferred and non-preferred saccade directions (*n* = 56 neurons, 4 mice). Shaded area, average ± s.e.m. Vertical orange bar, 0 - 90% rise time of saccades (26 ms). *P*-values, comparison of activity for preferred and non-preferred saccade directions in 20-ms bins (Wilcoxon signed-rank test, one-tailed; only significant values shown). Inset, magnified. (**C**) Left, schematic of pulvinar recording during saccades in TTX-blinded animals. Right, example neuron preferring temporal saccades. Raster plots (top), and PETH (bottom). (**D**) Left, scatter plot of the response to nasal and temporal saccades (for all responsive neurons). Right, average PETH of discriminating neurons (colored data points on left scatter plot) for preferred and non-preferred directions (*n* = 61 neurons, 12 mice). Baseline normalized. Shaded area, average ± s.e.m. Vertical orange bar, 0 - 90% rise time of saccades (26 ms). Inset, magnified. (**E**) Left, Schematic of pulvinar recordings during saccades in TTX-blinded animal and optogenetic antidromic activation of pulvinar projections to V1. Center, example of antidromically activated neuron. Raster plot (top) and spike probability (bottom; 0.05 ms bin). Blue shaded area indicates time of the 1-ms LED illumination. Right, response of neuron shown in center panel to saccades. This neuron prefers nasal saccades. Raster plot (top) and PETH (bottom). Dotted line in PETH, baseline firing rate. (**F**) Average PETH of saccade-responsive pulvinar neurons that were antidromically activated by illumination of V1 (*n* = 13 neurons, 3 mice). Shaded area, average ± s.e.m. Vertical orange bar, 0 - 90% rise time of saccades (26 ms). (**G**) Left, schematic of V1 recording during saccades in TTX-blinded mice before and after pulvinar silencing. Center, raster plot (top) and PETH (bottom) of an example neuron in response to nasal saccades before and after pulvinar silencing. Right, average PETH of saccade responsive neurons before and after pulvinar silencing (*n* = 56 neurons, 5 mice). All nasal and temporal saccades are included. Shaded area, average ± s.e.m. (**H**) LFP from example animal aligned to the time of saccades for layer 2/3 (L2/3), layer 5 (L5) and layer 6 (L6) before and after pulvinar silencing.

We next focused on the pulvinar, a higher-order thalamic nucleus with extensive projections to superficial layers of V1 (*27*), consistent with the estimated entry point of the non-visual input (see above), and a structure in which neurons have also been shown to respond to saccades (*26, 28, 29*). Indeed, recordings in the pulvinar in TTX-blinded animals revealed that almost half of the neurons responded to saccades (102 out of 225, 12 mice), many of which were also direction selective (61 out of 102; Fig. 4C, D). Furthermore, the PETH of directionally selective pulvinar neurons for the preferred and non-preferred directions diverged prior to saccade onset (200 ms window before onset, 0.7 ± 0.2 Hz evoked FR ± s.e.m. for the preferred direction, 0.2 ± 0.2 Hz for the non-preferred; Wilcoxon signed-rank test, one-tailed, *p* = 0.013, *n* = 61; Fig. 4D, right). We also verified the presence of direct projections from these neurons to V1, using Channelrhodopsin-2 mediated antidromic activation (*30–32*) (see Methods). Of 23 neurons that were identified in such a manner, more than half responded to saccades (13 out of 23, 3 mice), and 5 neurons discriminated saccade direction (Fig. 4E, F). To determine whether neurons in the pulvinar provide the non-visual saccadic input to V1, we silenced the pulvinar (using either TTX or muscimol), while recording from V1 in TTX-blinded animals (Fig. 4G). Strikingly, pulvinar silencing abolished both the non-visual saccadic response in V1 neurons (88 ± 19% average decrease ± s.e.m. in saccade evoked FR; Wilcoxon signed-rank test, one-tailed, *p* = 0.012, *n* = 56; based on 56 saccade responsive neurons out of 140 pre-silencing, 5 mice) and the saccade-triggered LFP (Fig. 4G, H, fig. S10). The silencing also strongly reduced the ability of V1 neurons to discriminate saccade directions (pre-silencing, 6.8 ± 1.6 Hz difference in evoked FR ± s.e.m. between preferred and non-preferred; post silencing, 1.3 ± 0.6 Hz, 74 ± 11% reduction; Wilcoxon signed-rank test, one-tailed, *p* < 0.0001, *n* = 29; based on 29 neurons that discriminated pre-silencing, 5 mice; fig. S10). Taken together, these results demonstrate that the pulvinar is the main source of the non-visual, saccade response in V1.

### The pulvinar input scrambles stimulus direction preference in V1

The anatomical separation between the source of the visual (dLGN) and the non-visual (pulvinar) inputs to V1 provides us with the experimental opportunity to silence the non-visual input while sparing the visual input. Silencing the pulvinar should lead to saccade responses in V1 that are mainly visual, driven by the shift of the image on the retina. In other words, pulvinar silencing should prevent scrambling, resulting in similar direction preference for real and pseudo saccades. We thus recorded the activity of V1 neurons in response to saccades made on the vertical grating, after injecting muscimol in the pulvinar, and compared it to the response to pseudo-saccades (Fig. 5A, B, fig. S11; distribution of saccade amplitudes was little affected by pulvinar silencing compared to control, fig. S2). Remarkably, with pulvinar silencing, direction selectivity for real saccades now predicted that for pseudo-saccades (Pearson ρ = 0.72, *p* < 0.0001, *n* = 29, 9 mice; Fig. 5C; a similar correlation was obtained also when including neurons lacking direction selectivity to pseudo-saccades; fig. S4). That is, the response of V1 neurons to a stimulus shifting on the retina was similar no matter whether the motion was induced externally or by a self-generated eye movement. Consistent with the correlation in the direction selectivity, the classifier trained on pseudo-saccades was able to decode the directions of saccade-induced visual motions at a level of performance similar to that for pseudo-saccades (74.7 ± 5.8% accuracy ± s.d. for pseudo-saccades with 250 neurons, chance level 46.8 ± 8.1%, Wilcoxon rank-sum test, two-tail, *p* < 0.0001, *n* = 300; 72.2 ± 5.9% for real saccades, chance level 54.2 ± 7.4%, Wilcoxon rank-sum test, two-tail, *p* < 0.0001, *n* = 300; Fig. 5D). Thus, the pulvinar provides a saccade-triggered, directionally selective non-visual input that precedes saccade onset and that, by combining with the visual input, scrambles direction selectivity for retinal image shifts in V1 (Fig. 5E).

**Fig. 5.**
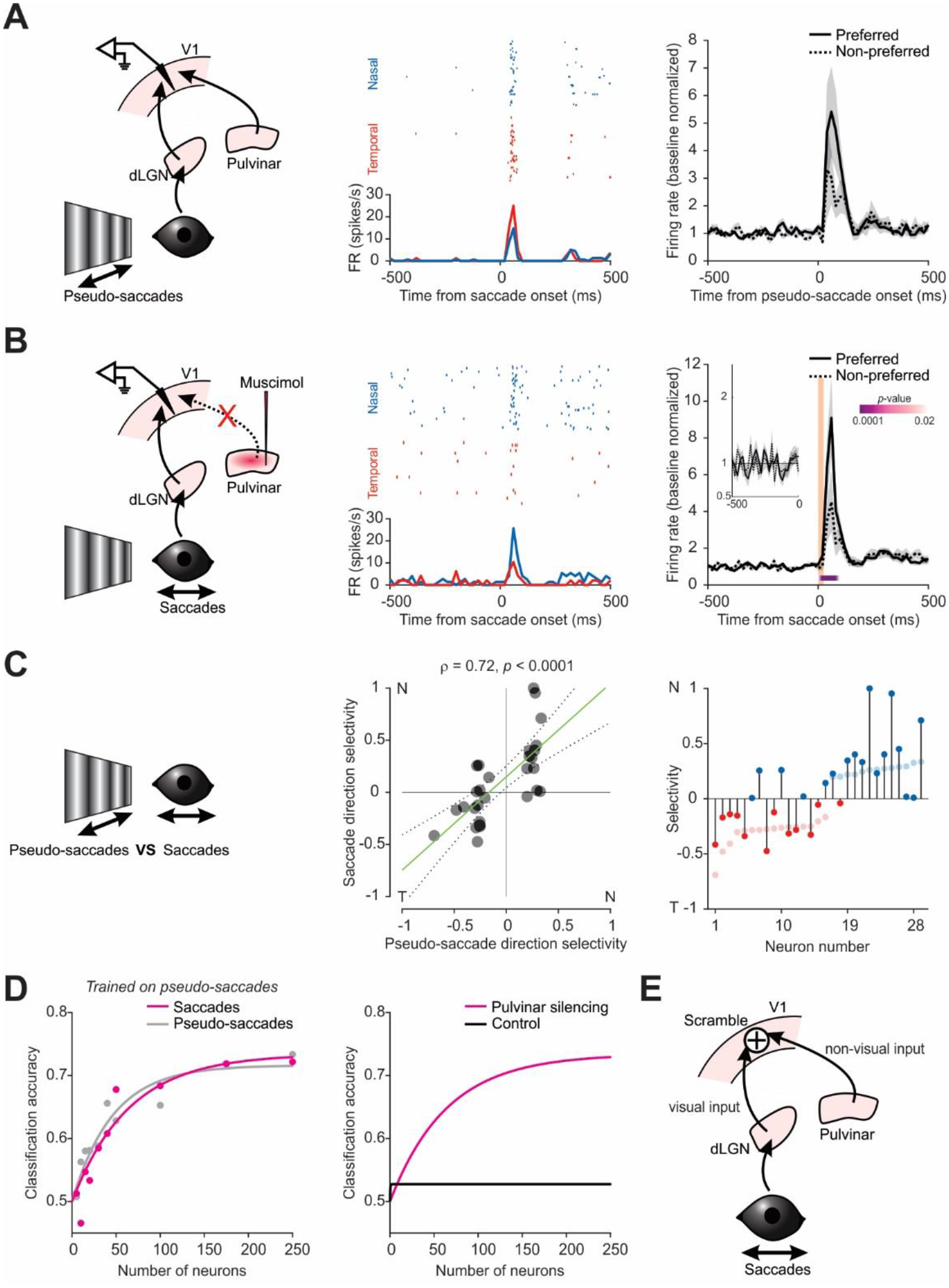
The pulvinar scrambles stimulus direction selectivity in V1 during saccades. (**A**) Left, schematic of V1 recording during pseudo-saccades. Center, example neuron preferring temporal saccades. Raster plots (top) and PETH (bottom). Right, average PETH of single neurons that discriminate nasal and temporal pseudo-saccades, sorted by their preferred and non-preferred pseudo-saccade directions (*n* = 29 neurons). Shaded area, average ± s.e.m. (**B**) Same as in (A), but for real saccade responses recorded after pulvinar silencing in the same animals. Center, example neuron preferring nasal saccades. Right, vertical orange bar, 0 - 90% rise time of saccades (25 ms). *P*-values, comparison of activity for preferred and non-preferred directions in 20-ms bins (Wilcoxon signed-rank test, one-tailed; only significant values shown). Note similar time-course of response to pseudo-saccades (A), including lack of separation prior to saccade onset (magnified in the inset). (**C**) Left, comparison of the response to saccades and pseudo-saccades. Center, scatter plot of direction selectivity for real and pseudo-saccades for all neurons discriminating pseudo-saccade direction. Green line, linear regression (coefficient 0.90, *p* < 0.0001); dotted lines, 95% confidence interval. Pearson ρ = 0.72, *p* < 0.0001, 29 neurons, 9 animals. Note good correlation, compared to conditions without pulvinar silencing (Fig. 2J). Right, neurons sorted according to their direction selectivity for pseudo-saccades (lighter shade), overlaid with direction selectivity for real saccades (darker shade). Note reduced scrambling compared to conditions without pulvinar silencing (Fig. 2K). (**D**) Left, classification accuracy of direction of motion (nasal or temporal) of a linear classifier trained on pseudo-saccades and tested on pseudo-saccades and real saccades, as a function of neurons included in the analysis. Lines, exponential fits. Note good accuracy on both pseudo- and real saccades. Right, classification accuracy for real saccades under control condition (black, from Fig. 2L) and under pulvinar silencing (magenta), for comparison. (**E**) The integration of saccade direction selective non-visual input from the pulvinar with saccade-induced visual motion results in scrambling of stimulus direction selectivity of V1 neurons during saccades.

## Discussion

Our study revealed that the response to external motion and saccades are distinct as early as in V1. This is because, during saccades, V1 combines a non-visual input that originates from the pulvinar and that depends on saccade direction with the visual input originating from the retina. The combination of non-visual and visual inputs scrambles stimulus direction preference of V1 neurons, due to the fact that the saccade direction preference of the non-visual input does not correlate with that of V1 neurons to visual input. As a consequence, the activity pattern in V1 in response to an image moving on the retina differs depending on whether that motion was generated by the motion in the environment or by eye movements.

The combination of two directionally selective yet uncorrelated inputs presents a simple yet effective strategy that enables mouse V1 to briefly change the representation of visual motion during saccades. Interestingly, similar changes in direction preference have been reported in higher visual areas of primates around the time of saccades, possibly indicating conserved neuronal mechanisms (*13, 33*). The change in representation of self-generated stimuli may work in concert with previously reported mechanisms in which sensory responses to self-generated stimuli are suppressed (*34–37*).

The dynamics of the V1 responses to saccades in freely moving and head-fixed animals were remarkably similar, warranting the use of head-fixed conditions to study the interaction between visual and non-visual inputs. Both in freely moving and head-fixed animals, V1 activity was modulated before saccade onset, peaked shortly after the saccade, outlasted the saccade duration and showed marked saccade direction preference. These dynamics likely reflect a motor command, reaching V1 via the pulvinar as an efference copy or corollary discharge. In support of this notion, the pulvinar has been postulated to receive an efference copy through collateral branches of cortical neurons that project to brainstem motor nuclei (*38–40*). Furthermore, the pulvinar also receives substantial input from the superior colliculus (*41–43*), a midbrain structure involved in saccade initiation (*44*). We cannot exclude, however, that later parts of the response may also reflect proprioceptive signals originating in the eye muscles. Since gaze shifts result from a coordination of eye and head movements (*5, 6*), the V1 response around the time of saccades may reflect the combined representation of eye and head motor commands. Whether V1 responses are dominated by eye motor command, as suggested by the similarity of the responses between freely moving and head-fixed conditions, or whether they also represent head motor command remains to be elucidated. Furthermore, while we have identified direct projections from saccade responsive pulvinar neurons to V1, other indirect routes also remain possible.

In humans, motion on the retina induced by saccades is often perceptually unnoticed, a phenomenon termed saccade suppression. If similar mechanisms to those revealed here are at work in the human brain, the distinct cortical representation of self and externally generated motion may prevent downstream areas from decoding the direction of visual motion induced by a subject’s own saccadic eye movement. Despite the lack of perceptual experience, however, studies in humans also show that visual processing remains active during saccades (*45, 46*). These results are consistent with our finding that the visual signal, rather than being suppressed, is combined with the non-visual input. Furthermore, our model suggests that the gain of the visual signal is reduced during saccades, consistent with the notion that a reduction in gain could further contribute to saccade suppression (*47*).

In conclusion, we have uncovered a circuit mechanism that allows V1 to distinguish motion induced by the animal’s own eye movement from changes in the environment through the combination of two independent inputs whose response properties are uncorrelated. This mechanism may represent a general strategy for sensory cortices to distinguish between self and externally generated stimuli.

## Materials and Methods

### Mouse handling

Experiments were conducted in accordance with the regulations of the Institutional Animal Care and Use Committee of the University of California, San Diego and of the University of California, San Francisco. All mice used in this study were wild-type C57BL/6J males or females from the Jackson Laboratory (JAX #000664) and between postnatal ages of three to six months.

Animals were familiarized to head-fixation for at least two weeks prior to recording. During this time, they were also familiarized to visual stimuli that would be used during recording. Animals were head-fixed on a custom-made passive treadmill, either circular or linear, and were free to run.

### Eye tracking

Video-oculography was used to track the movement of the right eye in both freely moving and head-fixed mice, contralateral to the hemisphere in which recordings were conducted.

In freely moving mice, the right eye was tracked using a miniature camera (Arducam Noir Spy Camera) mounted on a custom designed holder attached to the skull. The eye was illuminated using an infrared LED mounted on the holder. The video was acquired at 90Hz through Raspberry Pi 3B+ using RPiCamera-Plugin (*48*).

For head-fixed experiments, a high-speed camera (IMPERX, IPX-VGA-210-L) was fitted with a 45 mm extension tube, a 50 mm lens (Fujifilm, Fujinon HF50HA-1B), and an infrared (IR) pass filter (Edmund Optics, #65-796). Images were acquired at 200Hz through a frame grabber (National Instrument, PCIe-1427). An IR hot mirror (Edmund Optics, #43-958) was placed parallel to the antero-posterior (A-P) axis of the animal (1 inch from the eye) in between the animal and the LCD monitor, and the camera captured the image of the eye through its reflection. The camera was angled at 59° relative to the A-P axis. Three IR 880 nm LED emitters (Digi-Key, #PDI-E803) were used to illuminate the eye.

### Measuring angular position of the eye

Head-fixed animals: One of the three IR LEDs (see above) was aligned with the optical axis of the camera and served as a reference to calculate the pupil position. The pupil was identified by thresholding and fitting an ellipse. We computed α, the angular position of the eye, according to sin(α) = *d*/Rp, where *d* is the projected distance on the camera image between the center of the ellipse and the corneal reflection (CR) of the reference LED and where Rp is the length of the radius that connects the rotational center of the eye with the center of the pupil on the plane that harbors the pupil. Note that Rp is shorter than the radius of the eyeball. Rp was estimated prior to the experiments as follows: The camera, together with the reference LED, were swung by calibration angles γ of ± 10° along a circumference centered on the rotational center of the eye (more precisely, on the rotational center of the mirror image of the eye since the eye was imaged through an IR mirror) such that the CR of the reference LED remained stationary relative to the image frame of the camera. We used different values of *d* obtained with different γ to estimate Rp. Complicating the issue is the fact that Rp is not fixed but changes with the size of the pupil [i.e. the distance from the rotational center of the eye to the plane that harbors the pupil increases with the constriction of the pupil; (*49*)]. We thus computed Rp under various luminance conditions to change pupil diameter (Dp, i.e., the long-axis of the fitted ellipse) and obtained the following linear relationship: Rp = *r* - *a* * Dp, where *r* is the radius of the eyeball; *a* typically ranges between 0.05 and 0.25. During eye tracking in both freely moving and head-fixed animals, this relationship was used to determine Rp for every video frame based on the pupil diameter. In some mice, Rp was estimated using the relationship obtained from littermates or other similarly sized mice. The details of the eye-tracking method during head-fixation has been published previously (*50, 51*).

Freely moving mice: To delineate the pupil, eight points along the edge of the pupil were tracked post-hoc using DeepLabCut (*52*) and were fitted with an ellipse. The center of the pupil was defined as the center of the ellipse, the center of the projected eye on the camera C (equivalent to CR in head-fixed, see above) was estimated by using the orientations of the ellipses at multiple pupil positions, and *d* is the distance between the center of the eye and the center of the pupil. The angular position of the eye, α, was computed as in head-fixed animals according to sin(α) = *d*/Rp. Rp was estimated from the equation Rp = *r* – *a* * Dp obtained under head-fixation.

### Surgery

Mice were implanted with either a custom T-shaped head bar (head-fixed experiments) or three threaded screw inserts arranged in a triangle (head-fixed and freely moving experiments; McMaster-Carr, #92395A109). The implantation was done stereotactically using an inclinometer (Level Developments, DAS-30-R) connected to an USB I/O device (National Instruments, USB-6008), so that the axes of the electrode manipulators for acute, head-fixed recordings would be aligned to the A-P, medio-lateral, and dorso-ventral axes of the skull. Mice were anaesthetized with 1 - 1.5% isoflurane and kept on a feedback regulated heat pad to maintain body temperature at 37 °C (FHC 40-90-8D). Prior to surgery, mice were given buprenorphine subcutaneously. Prior to incision, topical lidocaine cream was applied to the skin. Once the scalp and fascia were removed, the head bar or the screw inserts were cemented using dental cement (Lang Dental, Ortho-Jet for head bars; 3M ESPE, Relyx Unicem2 for screw inserts). The animals were allowed to recover in their home cage for at least 1 week following the surgery.

For mice prepared for freely moving experiments, an extracellular electrode (Diagnostic Biochips, P64-4) mounted on a custom-designed hat for chronic recording was implanted one day prior to the recording session using dental cement. This procedure was performed weeks after the initial implantation of the screw inserts. Mice were anaesthetized with 1 – 1.5% isoflurane and kept on a feedback regulated heat pad. The electrode held by a holder was lowered to 1100 μm below pia surface using micromanipulators, and the hat was cemented in place before retracting the holder. The cranial window over V1 was ∼200 μm by ∼200 μm and covered with silicone gel after the electrode insertion to prevent V1 from drying. A ground wire (A-M Systems) was inserted in the cerebellum. A custom designed camera mount was also attached to the head using the previously implanted screw threads (see above).

In head-fixed experiments, cranial windows for extracellular recordings were made one or two days prior to the recording sessions. For all recordings, the size was ∼500 μm – 1 mm by ∼500 μm – 1 mm. Whiskers that would interfere with eye tracking were also trimmed at this point. Following the craniotomy, the window was sealed with biocompatible silicone sealant until the recording session (World Precision Instruments, Kwik-Cast).

The cranial windows were centered around the following coordinates that were marked during the head bar or screw insert implantation:

V1 recording: 2.7 mm lateral to the midline, 4.1 mm posterior to the Bregma Pulvinar recording: 1.2 mm lateral to the midline, 1.9 mm posterior to the Bregma dLGN recording: 2.4 mm lateral to the midline, 2.2 mm posterior to the Bregma For the identification of pulvinar neurons that send projections to V1 through optogenetic antidromic activation, AAV2/1.hSyn.ChR2(H134R)-eYFP.WPRE.hGH (Addgene 26973P) was injected in the pulvinar in the left hemisphere, prior to the implantation of the head bar or screw inserts.

### Visual stimulation

Visual stimuli were presented on an LCD monitor running at 240 Hz (Gigabyte, AORUS KD25F) to the right eye, contralateral to the hemisphere in which recordings were performed. The monitor was angled at 31° counterclockwise relative to the A-P axis of the animal and tilted 20° towards the animal relative to the gravitational axis. It was positioned such that the tangent point between the plane of the monitor and a sphere around the center of the eye was in the center of the monitor. The distance from the center of the eye to the tangent point was 133 mm, with the monitor covering 128° of the field of view horizontally and 97° vertically. In the experiment described in fig. S6 (a full-field flash), an LCD monitor running at 75 Hz was used.

The static vertical grating used in the experiments described in Fig. 2 and 5 was a full-field sinusoidal grating with 70% contrast, spatial frequency of 0.08 cycles per degree (cpd), and mean luminance of 40 to 60 cd/m^2^ (gamma corrected; a fixed luminance for each animal). It was spherically morphed around the center of the animal’s right eye to maintain the same spatial frequency across different spatial locations on the retina. For pseudo-saccades, the exact same grating was quickly shifted horizontally once every 1.5 seconds on average, over the span of 7 frames (6 inter-frame intervals, 25 ms). The speed of the shift over the 7 frames was linear. The direction and amplitude of each shift was pre-determined by randomly drawing from the distribution of real saccades collected separately from wildtype unmanipulated mice. For a nasal pseudo-saccade, the grating was shifted in the temporal direction, and for a temporal pseudo-saccade, the grating was shifted in the nasal direction. *Post-hoc*, every pseudo-saccade was checked for display errors such as a dropped frame. All pseudo-saccades that occurred within 500 ms of a real saccade were also discarded from further analyses, which resulted in about 350 pseudo-saccades for each animal over the span of 10 min. We then resampled the pseudo-saccades to match the direction and amplitude of the real saccades collected from the same animal. In order to increase the statistical power, we resampled two matching pseudo-saccade events for every saccade. The mean ± standard deviation of the difference in amplitude between a real saccade and its matched pseudo-saccades was 0.25 ± 0.45° (498 pseudo-saccades, 4 mice) for experiments in Figure 2 and 0.18 ± 0.45° (942 pseudo-saccades, 9 mice) in Figure 5.

For every animal, response to pseudo-saccades was collected at the beginning of the experiment. Response to real saccades using the static grating was collected after the pseudo-saccade session. The two responses were collected separately, in order to maximize our chances of obtaining saccades whose responses were not contaminated by pseudo-saccade responses.

To verify the absence of visual responses, following either intraocular TTX injection or muscimol injection in the dLGN, we used the following visual stimuli: For the intraocular TTX injections, we used full-field luminance change from 0 cd/m^2^ to 100 cd/m^2^ lasting 26 ms. For the muscimol injection in the dLGN, we used a full-field vertical grating (0.02 cpd; contrast 0.5), presented every 10 s for 32 ms, preceded and followed by a gray screen of the same average luminance of 40 cd/m^2^.

All visual stimulation protocols were custom written in LabVIEW (National Instruments) and MATLAB (Mathworks) using Psychophysics Toolbox 3 (*53–55*).

### Acute extracellular recording in head-fixed mice

All recordings in this study were performed on the left hemisphere. On the day of recording, animals were first head-fixed, and the Kwik-Cast sealant over the cranial windows was gently removed. Artificial cerebrospinal fluid (ACSF; 140 mM NaCl, 2.5 mM KCl, 2.5 mM CaCl_2_, 1.3 mM MgSO_4_, 1.0 mM NaH_2_PO_4_, 20 mM HEPES, and 11 mM glucose, adjusted to pH 7.4) was quickly applied to the craniotomy to prevent the exposed brain from drying. Different configurations of silicon probes were used over the course of the study: A2x32-5mm-25-200-177-A64 (NeuroNexus), A1x64-Poly2-6mm-23s-160-A64 (NeuroNexus), A1x32-Poly2-10mm-50s-177-A32 (NeuroNexus), and ASSY-77 H2 (Cambridge NeuroTech). Using a manipulator (Luigs & Neumann), the probes were slowly lowered to the recording site. For V1, probes were lowered to 1,000 μm from the pia; for the dLGN, 3,000 μm; and for the pulvinar, 2,900 μm. For recordings in the thalamus, the probes were painted with lipophilic DiI prior to the insertion visualization of the recording track. Successful targeting was verified *post-hoc*.

For optogenetic activation of the axon terminals of pulvinar neurons, a glass fiber-optic cable (960 μm core, NA = 0.63; Doric Lenses) connected to a 465 nm LED light source (Doric Lenses, #LEDC1-B_FC) was placed ∼500 μm above the craniotomy on V1. The light source was driven by an LED driver (Thorlabs, #LEDD1B) at 1000 mA for 1 ms every 6 s for 10 min (100 trials).

Recordings were started 15 minutes after the insertion of the probes. Signals were sampled at 30 kS/s using 64 channel headstages (Intan Technologies, #C3315) combined with adapters (NeuroNexus, Adpt.A64-Omnetics32_2x-sm), connected to an RHD USB interface board (Intan Technologies, #C3100). The interface board was also used to acquire signals from photodiodes (TAOS, #TSL253R) placed on the visual stimulation monitor as well as TTL pulses used to trigger the eye-tracking camera and the LED. These signals were used during analyses to synchronize visual stimulus timings, video acquisition timings, and LED photo-stimulation timings with electrophysiological recordings. All raw data were stored for offline analyses. Occasionally, we recorded from the same animal on two successive days, provided no pharmacological manipulation was performed on the first day. In these instances, the craniotomy was re-sealed with Kwik-Cast after the first recording session. For post-hoc histological analyses, brains were fixed in 4% paraformaldehyde (PFA) in phosphate buffer saline (PBS) overnight at 4 °C.

### Extracellular recording in freely moving mice

Mice were habituated in an acrylic open-air recording chamber under ambient light (length x width x height = 13.25 in. x 9 in. x 9.5 in.) for one hour each day for three days prior to the day of the recording. On the day of the recording, a miniature camera connected to Raspberry Pi (see Eye Tracking above) was mounted on the camera mount, and the implanted electrode was connected to an RHD USB interface board (Intan Technologies, #C3100). The TTL pulses from Raspberry Pi, used to synchronize the video frames with the electrophysiological signals, was also acquired through the interface board. Each recording session was 90 minutes long.

### Pharmacology

Intraocular injection of tetrodotoxin (TTX; 40 μM) was performed 2 hours prior to recording, under isoflurane anesthesia. A typical procedure lasted less than five minutes. Carbachol (0.011% w/v) was co-injected with TTX to prevent the pupil from fully dilating, since a fully dilated pupil reduces the accuracy of the eye tracking. Immediately prior to the injection, a drop of proparacaine hydrochloride ophthalmic solution was applied to the eye as a local anesthetic (Bausch + Lomb, 0.5%). TTX solution was injected intravitreally using a beveled glass micropipette (tip diameter ∼50 μm) on a micro injector (Nanoject II, Drummond) mounted on a manual manipulator. 1 μl was injected in each eye, at the speed of 46 nl/s. In some animals, the injection solution also contained NBQX (100 μM) and APV (100 μM). The animals were head-fixed for recording following a 2-hour recovery period in the home cage. The suppression of retinal activity was assessed for every experiment by the lack of response in visual cortex to a full-field flash of the LCD monitor.

Silencing of the dLGN and pulvinar was performed by injecting 30 nl of 5.5 mM muscimol-BODIPY at the speed of 300 nl/min, using a beveled glass pipette (tip diameter ∼20 - 40 μm) on a micro injector UMP3 with a Micro4 controller (World Precision Instruments). The injector was mounted on a micromanipulator (Luigs & Neumann) for stereotactic injection. In two of the pulvinar silencing experiments, TTX was used instead. The concentration of TTX was 60 μM, and 40 μl was injected at 40 μl/min. After the recording, brains were fixed in 4% PFA in PBS overnight at 4 °C for histological analysis of BODIPY on the next day.

### Histology

Anesthetized mice were perfused transcardially with 4% PFA in PBS (pH7.4). Brains were removed and further post-fixed in 4% PFA in PBS at 4 °C overnight, after which the solution was replaced with PBS. They were kept at 4 °C until they were coronally sectioned (100 μm sections) with a vibratome. Sections were mounted in Vectashield mounting media containing DAPI (Vector Laboratories H1500) and imaged with a camera (Olympus DP72) attached to an MVX10 (Olympus) stereoscope.

### Analyses

#### Detection of saccades

Head-fixed mice: Saccades were detected post-hoc from the eye tracking data, using a custom written algorithm in MATLAB. The algorithm searched for any event in which the angular position of the eye changed by more than 0.75 degrees along the horizontal axis in one video frame (5 ms). We discarded all events where the eye position did not move in the same direction for at least 3 successive frames (15 ms) and in which the peak amplitude of the eye movement was below 3 degrees. Furthermore, in order to eliminate the influence of preceding saccades on V1 responses, we only analyzed saccades that occurred in isolation, i.e., that were preceded by a period of at least 500 ms during which the eye did not move

Freely moving animals: The custom algorithm searched for events in which the eye position changed by more than 5.5 degrees in any direction in one video frame (11 ms). This equates to 500 degrees/s, exceeding the speed of most head movements in mice (*56*) and thus ensuring that the detected eye movements were not image stabilizing movements (i.e. vestibulo-ocular reflexes). The beginning of the saccade was defined as the first frame whose eye movement speed exceeded 200 degrees/s. The saccades were required to be at least two frames long (22 ms), and the vector of the eye movement for every frame of a saccade event was required to be within 45 degrees of each other.

#### Unit isolation

Single units from extracellular recordings were isolated using KiloSort (*57*) and visualized using Phy for further manual merging and splitting. The quality of isolated units was assessed using refractory period violations and stability of amplitude. The depth for each unit was assigned according to the electrode site at which its amplitude was the largest. For V1 recordings, units with trough-to-peak times longer than 0.5 ms were categorized as regular-spiking neurons. Units with shorter tough-to-peak times were categorized as fast-spiking neurons. Multi-units were defined as the collection of all units that remained after excluding noise using Phy. In the main text, we refer to isolated single units as neurons.

We used the spontaneous firing rate (FR) to register the recording depth across experiments. We approximated the border between layer 4 and layer 5 at ∼125 μm above the channel with maximum spontaneous FR. Channels within 200 μm below this border were assigned to layer 5 and channels within 150 μm above the border, to layer 4.

#### Inclusion criteria

Only animals with at least 15 saccades in each direction were analyzed. For this study, we focused on saccade-related activity of V1 neurons. Nonetheless, we found single units in our recordings whose activity correlated with the stationary eye position (putative “eye-position units”), both in control and TTX-blinded animals. Because there is a correlation between the direction of saccades and the position of the eye along the horizontal plane before the saccade (i.e., the more temporal the position of the eye before the saccade, the more likely the upcoming saccade will be nasal), some of these units were capable of discriminating the direction of future saccades, regardless of whether they responded to saccade onset. While these units represent a minority of the population, they would introduce a confounder in the current study because rather than discriminating saccade direction they code for eye position. Thus, for the analyses of single units in head-fixed animals, we excluded putative eye position units, i.e. units whose baseline activity (measured 500 ms prior to the onset of saccades) were significantly different between the two directions of the upcoming saccades (i.e. nasal and temporal). In freely moving experiments, all units were considered. The number of the excluded eye-position units, along with the total number of isolated single units for each experiment are detailed here:

Isolated single units from V1 recorded with a stationary grating, pseudo-saccades, or a gray screen without any manipulation (Fig. 2; Fig. 3A, B): 353 total single units (36 eye-position units), 4 mice.

Isolated single units from V1 recorded under TTX-blinding (Fig. 3C, D): 254 total single units (51 eye-position units), 8 mice.

Isolated single units from the dLGN recorded under TTX-blinding (Fig. 4A): 198 total single units (24 eye-position units), 4 mice.

Isolated single units from V1 recorded under dLGN silencing (Fig. 4B): 163 total single units (24 eye-position units), 4 mice.

Isolated single units from the pulvinar recorded under TTX-blinding (Fig. 4C, D): 261 total single units (36 eye-position units), 12 mice.

Isolated single units from V1 recorded under TTX-blinding and pulvinar silencing (Fig. 4G): 156 total single units (16 eye-position units, prior to pulvinar silencing), 5 mice.

Isolated single units from V1 recorded with a stationary grating with pulvinar silencing (Fig. 5A, B): 328 total single units (32 eye-position units), 9 mice.

#### Response to saccades and pseudo-saccades

Saccades in freely moving animals were categorized into eight evenly spaced directions. To determine whether a unit is responsive to saccades, we proceeded as follows: We performed Kruskal-Wallis test using its response and its baseline activity in each of the eight directions (total 16 categories). The response was defined as the number of spikes within 100 ms from the onset of saccades, while baseline activity was defined as the number of spikes in a 100 ms window between −300 ms to −200 ms from the onset. If the unit passed this test (critical value 0.05), we proceeded to perform multiple comparisons among the 16 categories using Tukey’s honestly significant difference procedure. The unit was considered responsive if the average response to any of the eight directions was 50% above or below the average baseline activity for the corresponding direction and met at least one of the following two criteria: 1) there was a significant difference between the baseline and the response for at least one direction; 2) there was a significant difference between the responses to any two of the eight directions.

In head-fixed experiments, units were considered responsive to saccades if they met either one of the following two criteria: If the number of spikes elicited within 100 ms after saccade onset was significantly different from the baseline for either nasal or temporal direction (the baseline was calculated as the number of spikes within a 100 ms window between −300 ms to −200 ms from the onset); Or if the number of spikes elicited within 100 ms after saccade onset were significantly different between nasal and temporal directions. Statistical significance was determined by rank-sum test, with a critical value of 0.05.

All reported responses in the main text are average FRs within the 100 ms window after saccade onset, unless otherwise noted.

#### Direction selectivity and discriminability

Direction selectivity of each single unit was calculated as the area under the receiver operating characteristics curve (AROC), linearly rescaled to the range from −1 to 1 (Gini coefficient). That is, 2*AROC – 1. In head-fixed animals, the selectivity was calculated based on two directions, nasal and temporal. The order was fixed, such that negative values indicate a preference for temporal saccades, and positive values indicate a preference for nasal saccades, i.e., the sign of direction selectivity corresponds to the preferred direction. To calculate the selectivity, the total number of spikes within the first 100 ms after saccade or pseudo-saccade onset from each event was used, without baseline subtraction. In freely moving animals, the preferred direction was defined as the direction with the maximum average firing rate within the first 100 ms after saccade onset. The selectivity was calculated based on the preferred direction and the non-preferred direction (direction opposite to the preferred). Discriminability was defined as the absolute value of the direction selectivity. Statistical significance of discriminability was calculated using rank-sum test with a critical value of 0.05. The direction selectivity index (fig. S1) was defined as (R_pref_ - R_nonpref_) / (R_pref_ + R_nonpref_), where R_pref_ and R_nonpref_ are the number of spikes within the first 100 ms after saccade onset in the preferred and non-preferred direction, respectively.

#### Average PETH with baseline normalization

When generating average PETHs with baseline normalization, neurons with baseline below 0.5 Hz were excluded to avoid substantial biases resulting from extremely low FR. The baseline of each neuron to saccades or pseudo-saccades was calculated using its mean activity 500 ms to 200 ms before the onset. For other visual stimuli, such as a full-field flash, mean activity between −200 to 0 ms before the onset was used. Note that this process was applied for *visualization purposes only*, and all statistics such as the calculation of direction selectivity, discriminability, direction selectivity index and the differences in evoked FRs were performed on all relevant neurons. Below, we report, for each figure panel, the number of relevant neurons and, in parenthesis, the number of neurons with baseline < 0.5 Hz: Fig. 1H: 53 (3); Fig. 2F, right: 97 (9); Fig. 2H, right: 44 (11); Fig. 3B, right: 109 (7); Fig. 3D, right: 64 (4); Fig. 4A: 49 (8); Fig. 4B: 56 (9); Fig. 4D, right: 61 (1); Fig. 4F: 13 (0); Fig. 4G, right, pre-silencing: 56 (6); Fig. 4G, right, post-silencing: 56 (8); Fig. 4A, center: 135 (12); Fig. 5A, right: 53 (4); Fig. 5B, center: 102 (10); Fig. 5B, right: 29 (3); fig. S9B: 29 (6). Statistical significance of the difference between PETHs for the preferred and the non-preferred direction was calculated for each 20 ms bin. This was calculated by signed-rank test, and the statistical significance was determined by adjusting the critical value (0.05) through Benjamini-Hochberg procedure to control the false discovery rate.

#### Modeling of saccade response on vertical grating with visual and non-visual inputs

Saccade responses on vertical grating (the number of evoked spikes within 100 ms from saccade onset) were predicted from 1) pseudo-saccade response, 2) saccade response on gray screen, or 3) the sum of the two responses. All responses were baseline subtracted values. The model is a linear regression (five-fold cross validated) with no intercept, followed by thresholding which ensured that the predicted FR did not fall below 0 Hz. That is, if the predicted decrease in evoked number of spikes exceeded the baseline FR, the value was adjusted so that the sum of the prediction and the baseline was zero. The explained variance is calculated as the Explained Sum of Squares divided by the Total Sum of Squares.

#### Identification of pulvinar neurons with axonal projections to V1 through antidromic activation

V1 was illuminated with 1-ms long pulses (100 trials) of 465 nm blue LED to induce antidromic spikes (see above). Success of antidromic activation was defined by two criteria: 1) greater than 20% probability of observing at least one spike within 5 ms from the onset of LED across trials and 2) less than 0.5 ms jitter (i.e. the standard deviation of the latency distribution of the first spikes occurring within the 5 ms window from LED onset was less than 0.5 ms).

#### Classification of saccade direction in head-fixed mice

We classified the direction of saccades and pseudo-saccades using linear discriminant analysis (LDA) and logistic regression on the number of spikes elicited by each single unit within 100 ms from the onset of each event (baseline subtracted). The procedure was preceded by principal component analysis (PCA) for dimensionality reduction. Only single units with average *fr* above 0.5 Hz was used. For each event of saccades or pseudo-saccades, the classifier assigned either nasal or temporal direction.

Training data consisted of the response to selected pseudo-saccades. This set of pseudo-saccades was selected such that the amplitudes and the number of events for nasal and temporal directions were matched. This ensured that the classifier depended on direction selectivity of each unit, rather than the difference in pseudo-saccade amplitudes or frequencies. The training data set was first subjected to PCA. Using the first dimensions that cumulatively accounted for 70% of total explained variance, we then trained LDA and logistic regression for classification. The resulting models for PCA, LDA, and regression were applied to the test data set, which were either responses to real saccades or pseudo-saccades whose amplitudes and directions had been matched to the real saccades.

As the same set of pseudo-saccades was presented to each animal during the recording (about 350 per animal, see Methods section on visual stimulation), we pooled data from single units across different animals for the same pseudo-saccade for the training data set. On the other hand, for the test data set, saccade data or the matched pseudo-saccade data from different animals were pooled simply based on the direction, and the amplitude was not considered. This is due to the difficulty in obtaining saccades of matching amplitudes from multiple animals. To mitigate the performance biases resulting from pooling combinations, 20 different combinations were generated. This pooling procedure potentially underestimates the performance of the classifier. Nonetheless, our focus was the comparison of the performance on saccades versus pseudo-saccades, both of which were subjected to the same procedure.

In order to calculate the classifier performance as a function of the number of single units used for the classification, a random subset of units (5, 10, 15, 20, 30, 40, 50, 100, 175, or 250 units) was chosen from the pooled data without replacement, before being subjected to training and testing. The entire procedure of the classification, including the random selection of units, was repeated 15 times for each number of units for every pooling combination (20, see above), and the average performance was calculated (i.e., average of 15 x 20 = 300 results). An exponential function, forced to go through 50% accuracy with 0 units, was fit to the data points for visualization. Chance levels were calculated by shuffling the evoked number of spikes relative to the saccade direction in the test data set.

## Acknowledgements

We thank L. Bao, J. Evora, N. Kim, Y. Li, M. Mukundan, and B. Wong for technical assistance; Riccardo Beltramo, E.J. Chichilnisky, Baohua Liu, Roger Nicoll, and Sarah Ruediger for critical reading of the manuscript; and all the members of the Scanziani lab for discussion.

## Author contributions

S.K.M. and M.S. designed the study. S.K.M. conducted all experiments and analyses. S.K.M. and M.S. wrote the manuscript.

## Data and materials availability

Datasets supporting the findings and custom codes used in this paper are available from the corresponding authors upon reasonable request.

---

**Fig. S1.**
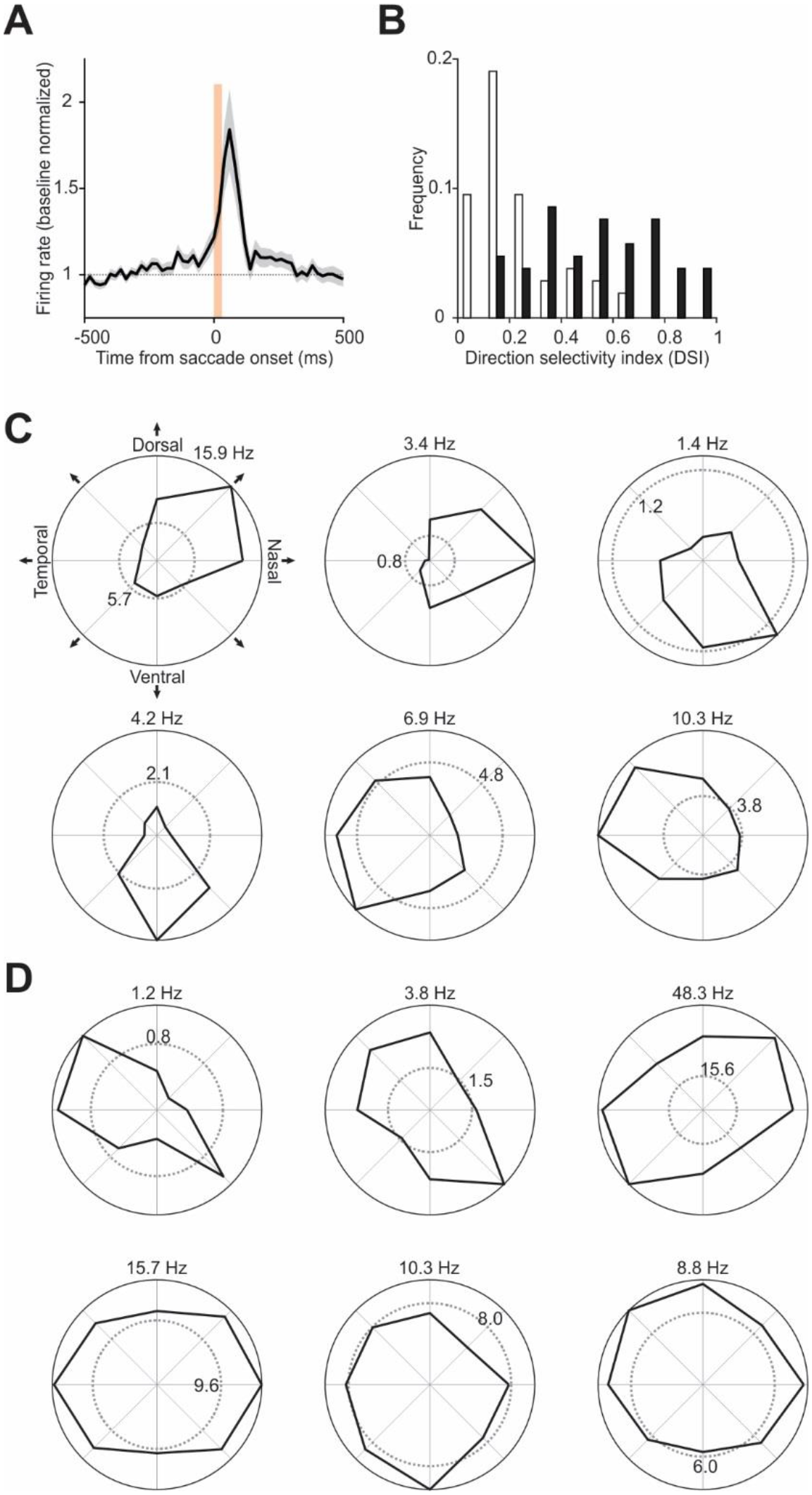
Response of V1 neurons to saccades in freely moving mice. (**A**) Average PETH of saccade responsive neurons (*n* = 105 neurons, 5 mice). For each neuron, the saccade direction with maximum response was taken. Baseline normalized. Shaded area, average ± s.e.m. Vertical orange bar, 0 – 90% rise time of saccades (31 ms). (B) Histogram of classical direction selectivity index (DSI; see Methods). White, non-direction selective (*n* = 52 neurons); black, saccade direction selective (*n* = 53 neurons). Significance was calculated by rank sum test (critical value 0.05). (**C**) Polar plots of example direction selective neurons. Black, saccade response; dotted gray, baseline. Firing rate for the preferred direction for each neuron is shown at the top. Numbers along the baseline indicate baseline firing rate, averaged for all directions. (**D**) Same as (C), but for non-direction selective neurons.

**Fig. S2.**
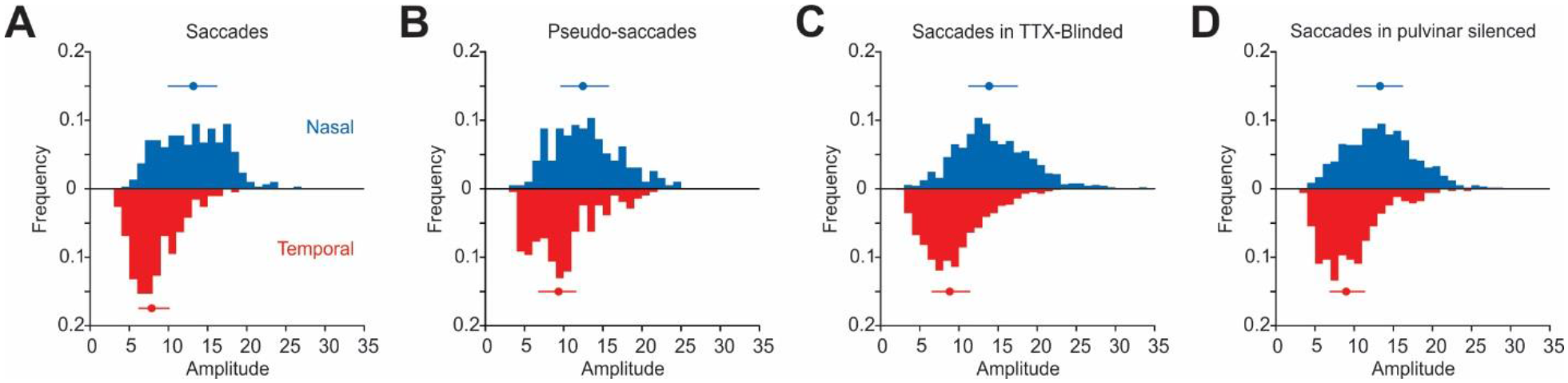
Distribution of real and pseudo-saccade amplitudes. (**A**) Histogram of saccade amplitudes in control mice. Blue, nasal saccades; red, temporal saccades. Bars indicate 25% and 75% quartile range, dot denotes median. Nasal saccades: median, 13.2°; 25% quartile, 9.9°; 75% quartile, 16.2°; *n* = 296. Temporal saccades: median 7.9°; 25% quartile, 6.2°; 75% quartile, 10.26°; temporal, *n* = 189. 4 mice. (**B**) Histogram of pseudo-saccade amplitudes. For analyses, pseudo-saccades with amplitudes matched to real saccades were further selected for each animal. All animals were presented with the same pseudo-saccades. Nasal pseudo-saccades: median, 12.5°; 25% quartile, 9.6°; 75% quartile, 15.8°; *n* = 193. Temporal pseudo-saccades: median 9.4°; 25% quartile, 6.7°; 75% quartile, 11.6°; *n* = 207. (**C**) Same as in (A) but for TTX-blinded animals. Nasal saccades: median, 13.9°; 25% quartile, 11.2°; 75% quartile, 17.5°; *n* = 887. Temporal saccades: median 8.8°; 25% quartile, 6.5°; 75% quartile, 11.5°; temporal, *n* = 561. 8 mice. (**D**) Same as in (A) but for animals with the pulvinar silenced. °Nasal saccades: median, 13.3°; 25% quartile, 10.4°; 75% quartile, 16.3°; *n* = 579. Temporal saccades: median 9.0°; 25% quartile, 6.9°; 75% quartile, 11.4°; temporal, *n* = 329. 9 mice.

**Fig. S3.**
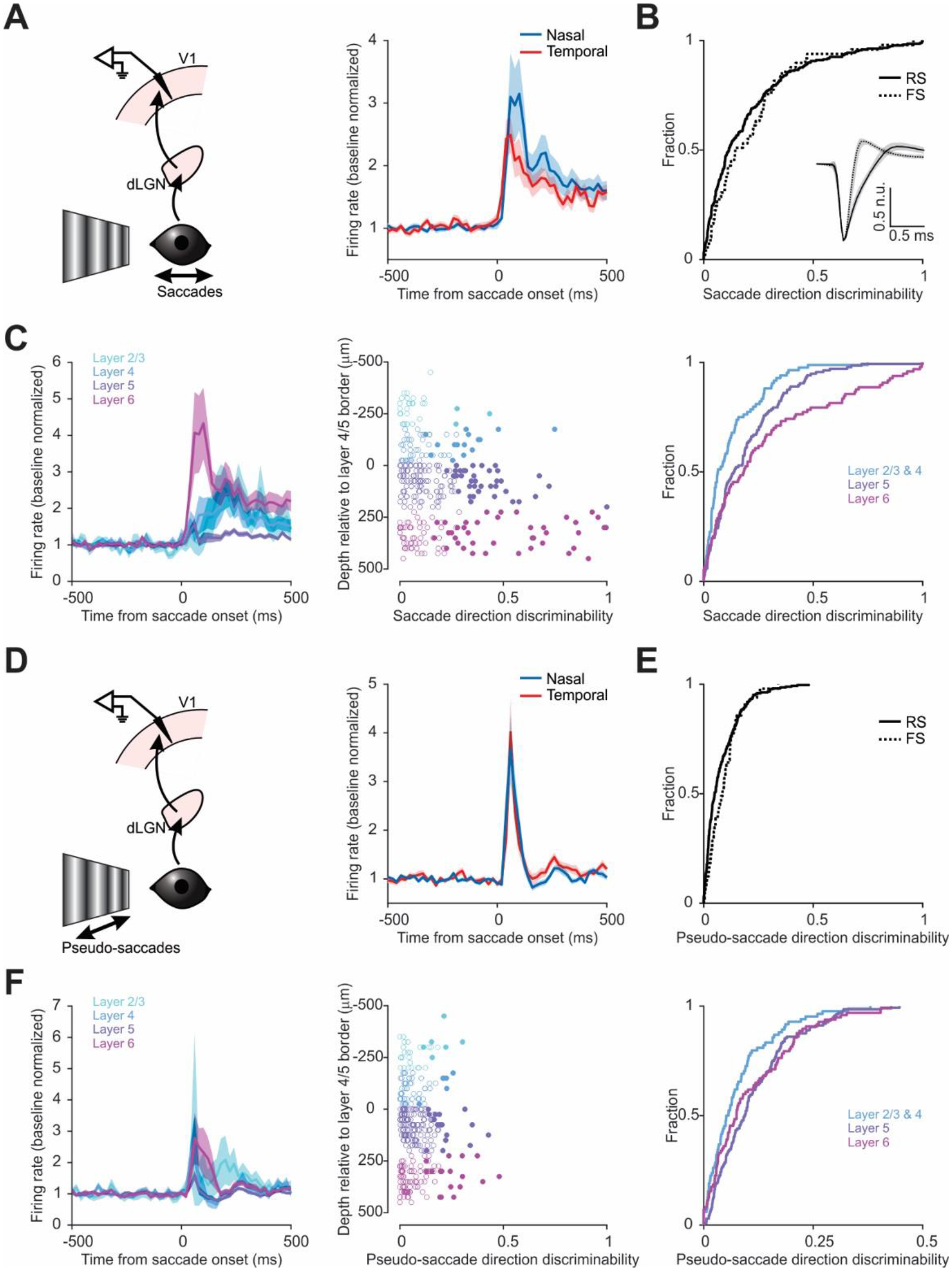
Dependence of saccade and pseudo-saccade response on spike waveform and cortical depth. (**A**) Left, schematic of V1 recording during saccades on a vertical grating. Right, average PETH of saccade responsive neurons to nasal and temporal saccades. Baseline normalized. Shaded area, average ± s.e.m. (**B**) Cumulative frequency distribution (CFD) of direction discriminability (absolute value of direction selectivity, see Methods), plotted for all regular-spiking (RS) and fast-spiking (FS) neurons. Anderson-Darling test, *p* = 0.28, *n* = 268 for RS, 49 for FS, 4 mice. Inset, median spike shape normalized to the trough. Shaded area, 25% and 75% quartiles. (**C**) Left, average PETH of all neurons to saccades according to cortical depth. All nasal and temporal saccades are included. *n* = 30 for layer 2/3, 52 for layer 4, 135 for layer 5, and 95 for layer 6. Baseline normalized. Shaded area, average ± s.e.m. Center, scatter plot of direction discriminability (x-axis) of all units as a function of cortical depth (y-axis). Open circles, statistically non-significant; filled, significant (cutoff *p* < 0.05, Wilcoxon rank-sum test). Color code as in left. Right, CFD of discriminability by cortical depth. Layers 2/3 and 4 were grouped together. 3-sample Anderson-Darling test, *p* < 0.0001. (**D-F**) Same as (A-C) but for pseudo-saccade responses in the same neurons. Note *n* is the same as (A-C). (E) Anderson-Darling test, *p* = 0.065. (F) Right, 3-sample Anderson-Darling test, *p* = 0.12.

**Fig. S4.**
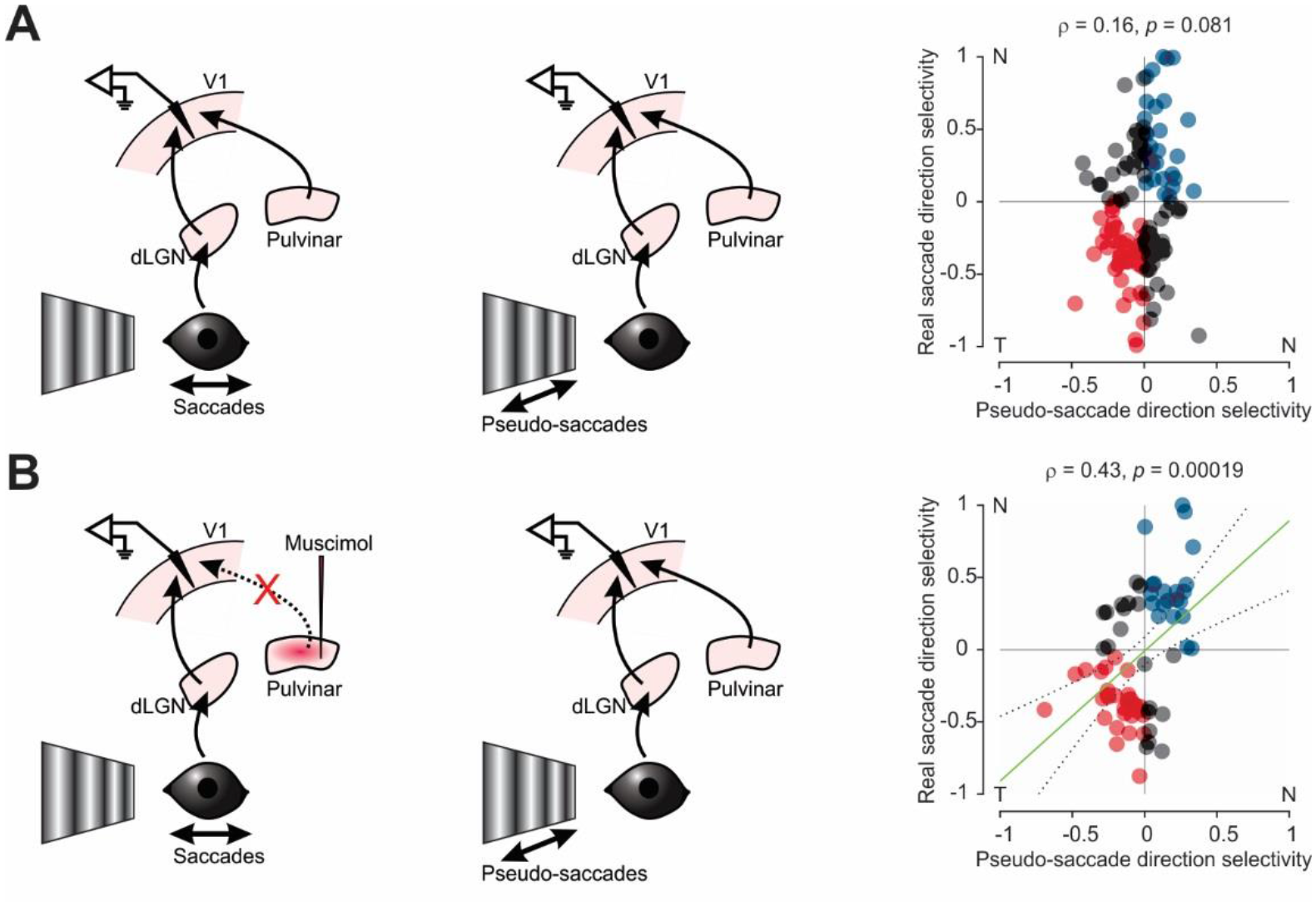
Direction selectivity for saccade-inducted motion and pseudo-saccades become correlated upon pulvinar silencing. (**A**) Comparison of the response to saccades and pseudo-saccades in control. Right, scatter plot of direction selectivity for real and pseudo-saccades for all neurons discriminating pseudo-saccade directions and/or real saccade directions. Blue, prefer nasal for both real and pseudo-saccades; red, prefer temporal for both real and pseudo-saccades; gray, prefer different direction for real and pseudo-saccades. Note the lack of a significant correlation (Pearson ρ = 0.16, *p* = 0.081, *n* = 127, 4 mice). (**B**) Same as in (A), but saccade response was recorded after pulvinar silencing. Right, direction selectivity became correlated (Pearson ρ = 0.43, *p* = 0.00019, *n* = 70, 8 mice). Green line, linear regression (coefficient 0.90, *p* = 0.0002); dotted lines, 95% confidence interval.

**Fig. S5.**
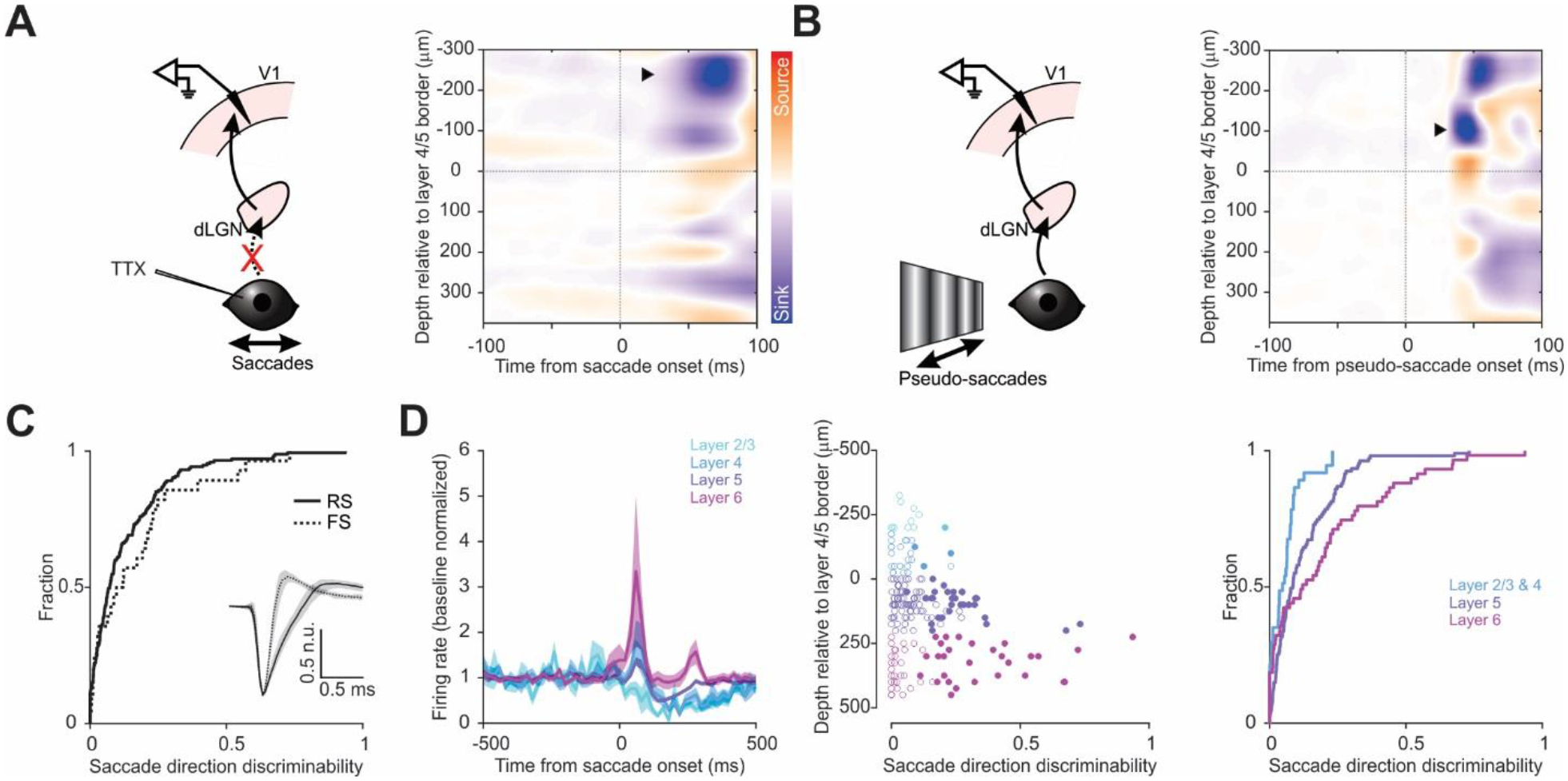
Dependence of saccade response on spike waveforms and cortical depth in TTX-blinded mice. (**A**) Left, schematic of V1 recording during saccades in TTX-blinded animals. Right, heatmap of the current source density (CSD) analysis from an example animal, showing a major sink in the supragranular layers (arrowhead). Data from 89 nasal and 82 temporal saccades. (**B**) Same as in (A), but for pseudo-saccades in a control animal. Note early sink in layer 4 (arrowhead). Color scale same as in (A). (**C**) CFD of saccade direction discriminability for RS and FS neurons in TTX-blinded animals. Anderson-Darling test, *p* = 0.090, *n* = 175 for RS, 28 for FS. Inset, median spike shape normalized to the trough. Shaded area, 25% and 75% quartiles. (**D**) Left, average PETH of all neurons to saccades in TTX-blinded animals. All nasal and temporal saccades are included. Layer 2/3, 13 neurons; layer 4, 24; layer 5, 107; layer 6, 59. Shaded area, average ± s.e.m. Center, scatter plot of direction discriminability of all units (x-axis) as a function of cortical depth (y-axis). Open circles, statistically non-significant; filled, significant (critical value 0.05, Wilcoxon rank-sum test). Color code as in left. Right, CFD of discriminability by cortical depth. Layers 2/3 and 4 were grouped together. 3-sample Anderson-Darling test, *p* < 0.0001.

**Fig. S6.**
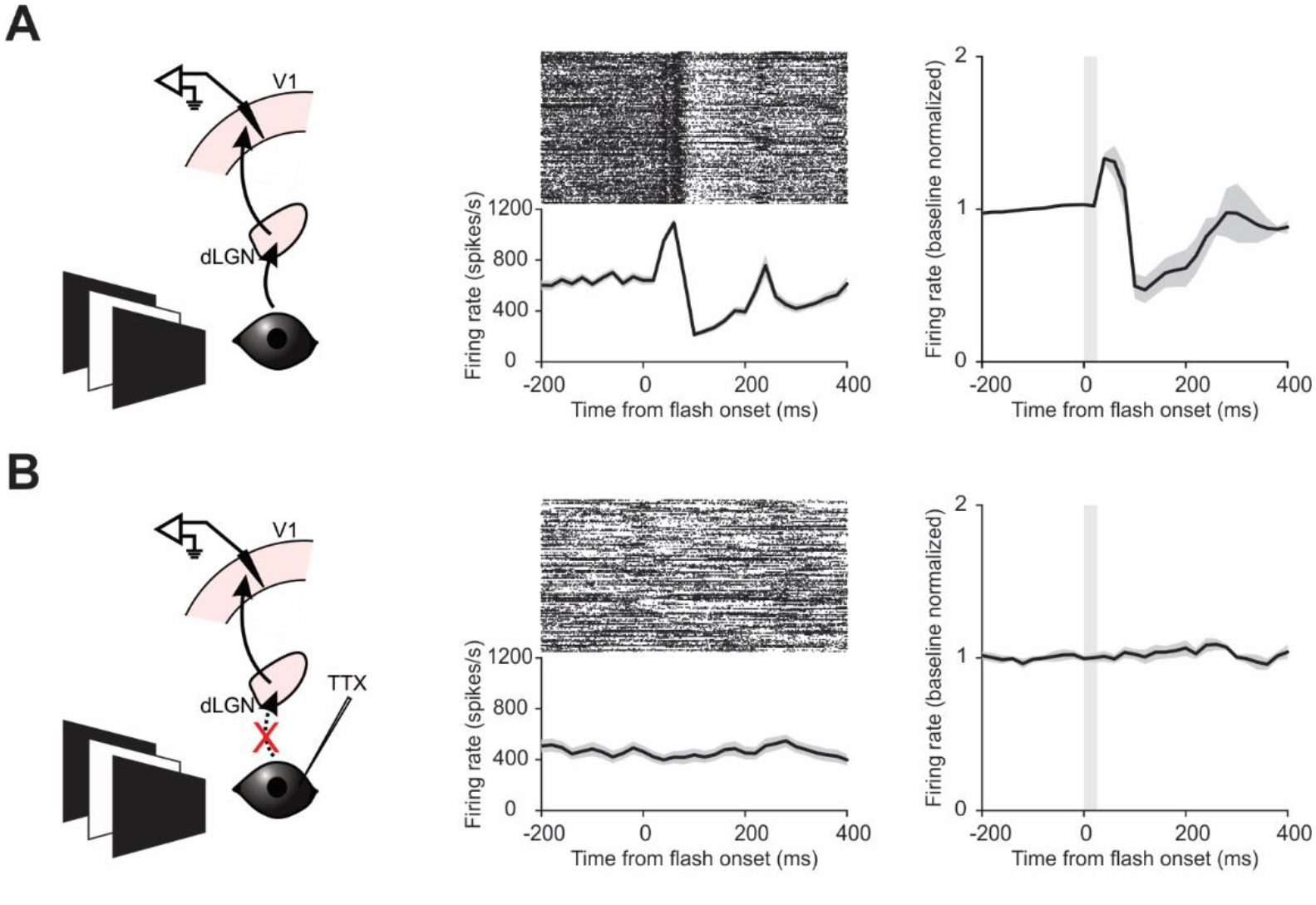
Intraocular injection of TTX abolishes visual response in V1. (**A**) Left, schematic of V1 recording during a brief (26 ms) full-field flash. Center, multi-unit response from an example recording. Raster plot (top) and average PETH (bottom). Right, average PETH of 4 mice. Baseline normalized. Shaded area, average ± s.e.m. 137.9 ± 31.2 Hz (22.1 ± 4.3%) evoked FR ± s.e.m. in a 60 ms window post onset (from +10 ms post-onset to +70 ms). (**B**) Same as in (A), but in TTX-blinded animals (8 mice). Note the lack of visual response. −8.0 ± 15.4 Hz (0.12 ± 3%) evoked FR ± s.e.m. in the 60 ms window post onset. Wilcoxon rank-sum test, one-tail, *p* = 0.0020.

**Fig. S7.**
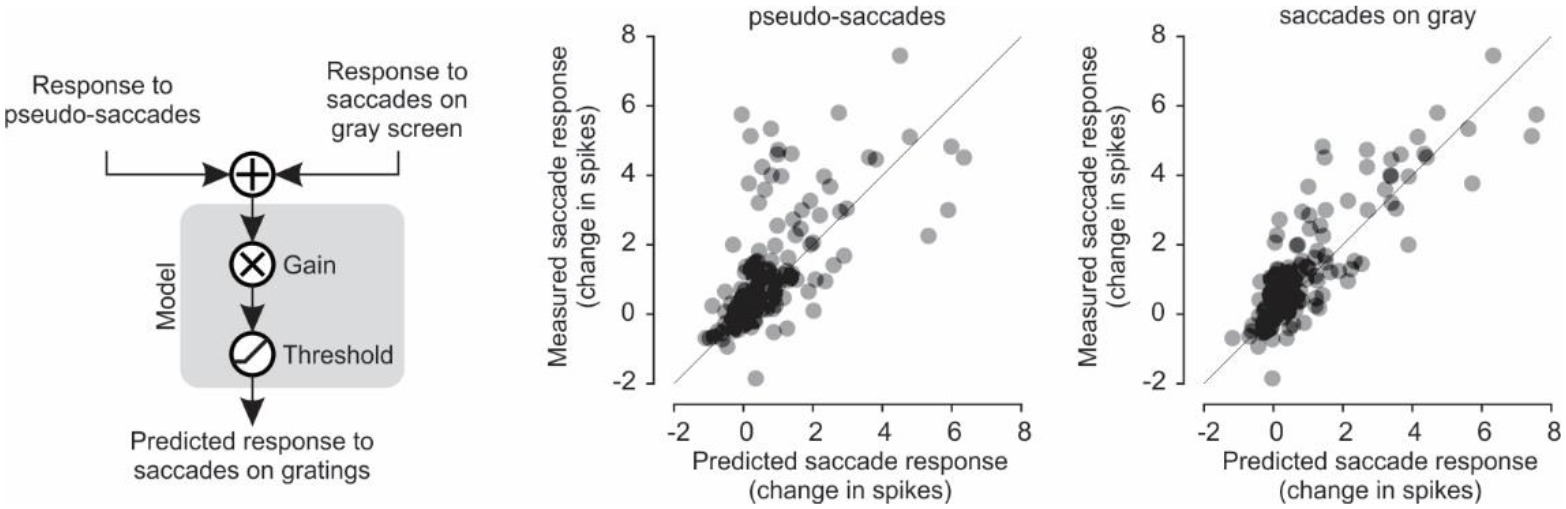
Modeling saccade response on a vertical grating with response to visual or non-visual input. Left, schematic of the linear regression-based model used to predict the number of spikes evoked by saccades on a vertical grating. The model is based on the response of neurons to pseudo-saccades and to saccades on a gray screen. Results from the sum of the two inputs are shown in Fig. 3G, in which the model explains 83% of the observed variance. Center, predicted number of spikes from the response to pseudo-saccades alone (x-axis) plotted against the observed values (y-axis). This model explains only 32% of the observed variance (gain 0.98, *p* < 0.0001). Right, predicted number of spikes from the response to saccades on a gray screen alone (x-axis) plotted against the observed values (y-axis). This model explains only 69% of the observed variance (gain 0.89, *p* < 0.0001).

**Fig. S8.**
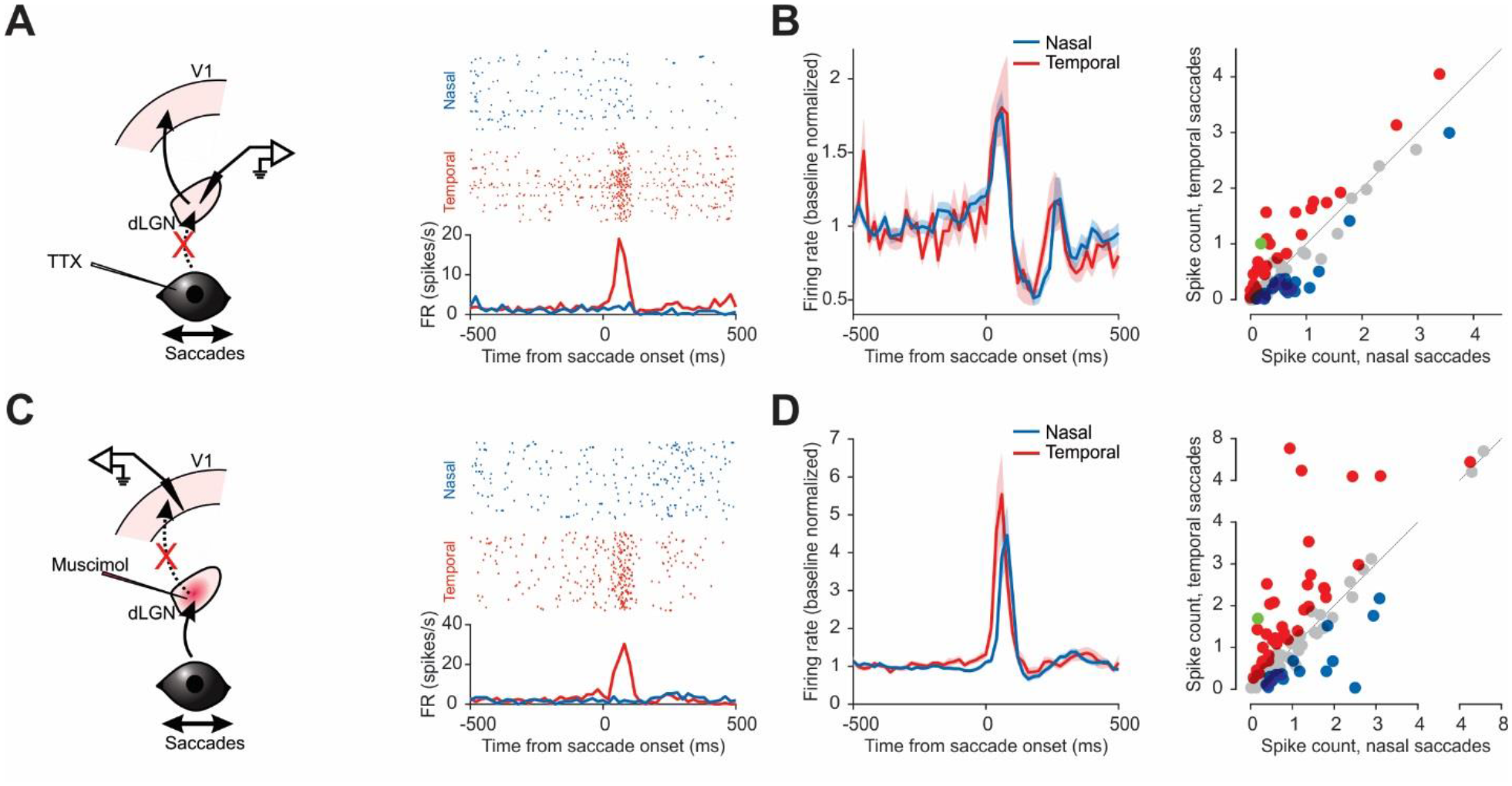
The dLGN is not the source of non-visual saccade response in V1. (**A**) Left, schematic of dLGN recording during saccades in TTX-blinded animals. Right, example neuron preferring temporal saccades. Raster plots (top) and PETH (bottom). (**B**) Left, average PETH of saccade responsive neurons to nasal and temporal saccades. *n* = 83, 4 mice. Baseline normalized. Shaded area, average ± s.e.m. Right, scatter plot of the response to nasal and temporal saccades (average spike count in a 100 ms window from saccade onset), for all responsive neurons. Blue, prefer nasal saccades; red, prefer temporal saccades; gray, no statistical difference; green, example neuron in (A). (**C-D**) Same as in (A-B), but for V1 neurons under dLGN silencing in TTX-blinded animals. (D) Left, *n* = 106, 4 mice. Right, Note direction selective neurons in V1 under dLGN silencing.

**Fig. S9.**
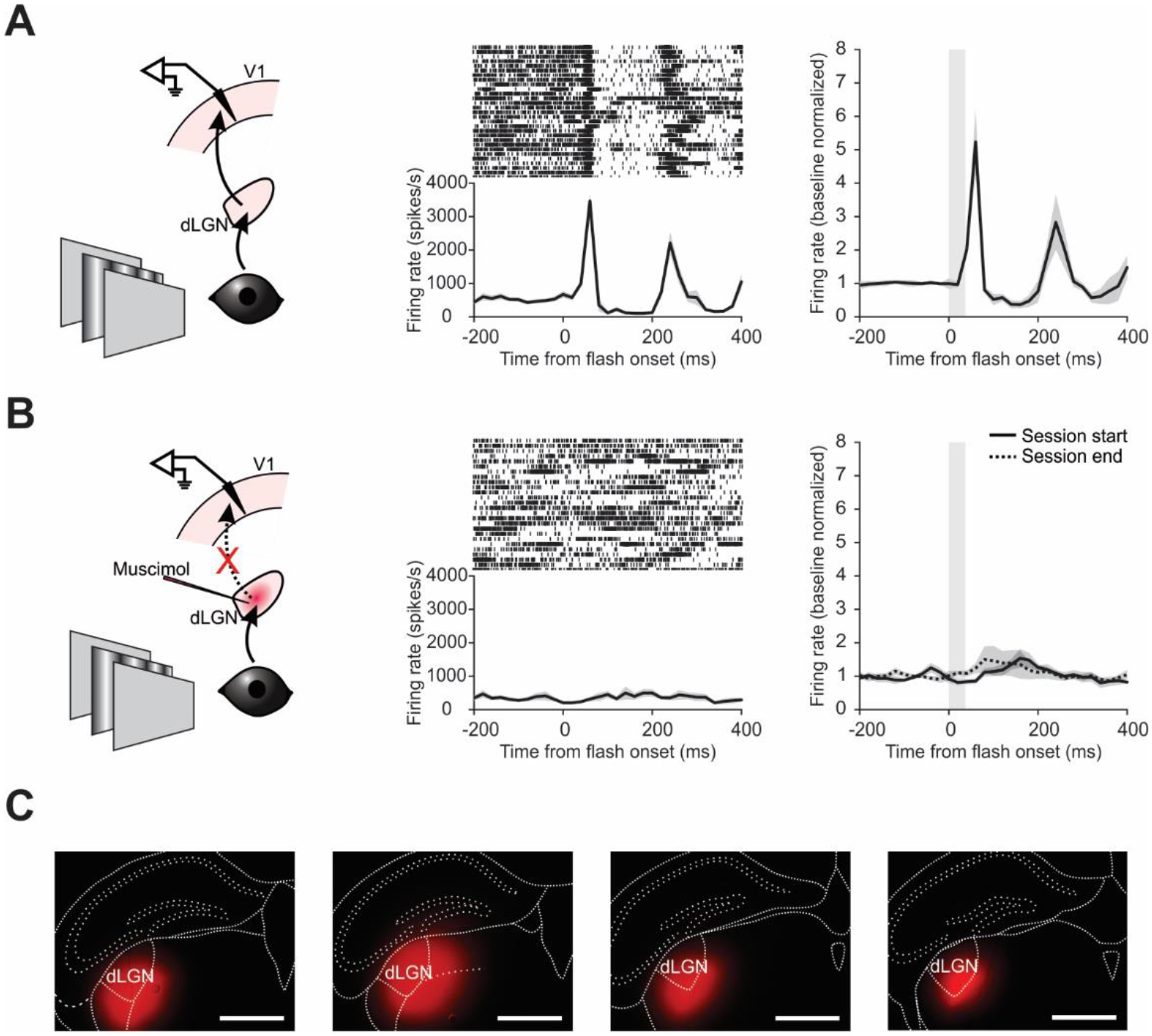
Muscimol-BODIPY injection in the dLGN blocks visual response in V1. (**A**) Left, schematic of V1 recording during a brief (32 ms) presentation of a full-field grating (see Methods). Center, multi-unit response from an example recording. Raster plot (top) and PETH (bottom). Right, average PETH of 3 mice. Baseline normalized. Shaded area, average ± s.e.m. 173.5 ± 29% average increase in evoked FR ± s.e.m. (**B**) Left and center, same as in (A), but for mice injected with muscimol-BODIPY in the dLGN. Right, average PETH of 4 mice. Recording started after muscimol injection. Response was measured both at the start of the recording session and at the end. Note the lack of visual response in both cases. Baseline normalized. Shaded area, average ± s.e.m. Average increase in evoked FR ± s.e.m. was −16.7 ± 7.5% (start) and 14.1 ± 8.8% (end). (**C**) Section images of the four mice injected with muscimol-BODIPY in the dLGN in (B). Red, BODIPY. Scale bar, 1 mm.

**Fig. S10.**
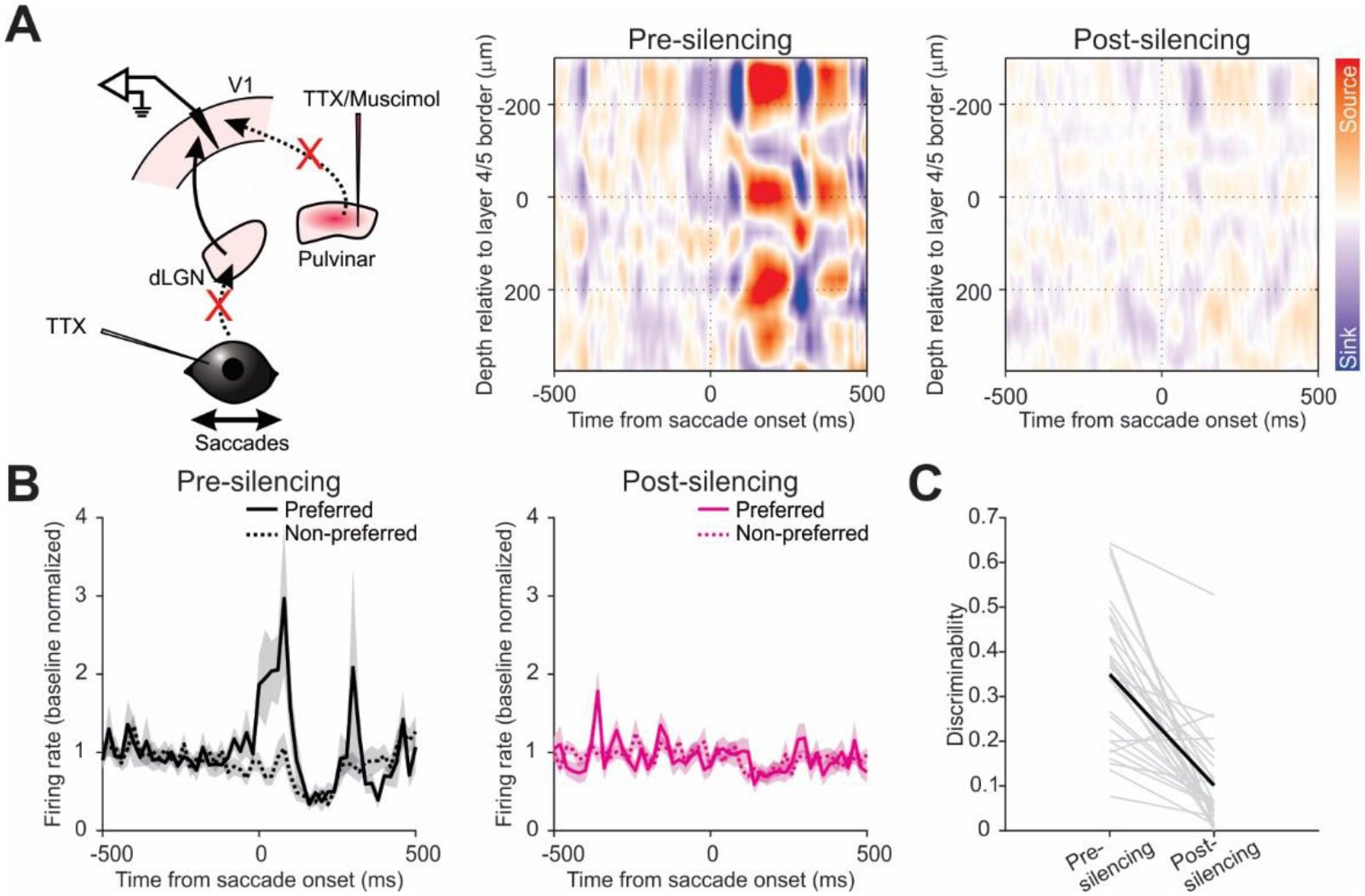
Silencing the pulvinar eliminates non-visual saccade input in V1. (**A**) Left, schematic of V1 recording during saccades in TTX-blinded animals before and after pulvinar silencing. Center, heatmap of the CSD analysis of an example animal, prior to pulvinar silencing. All nasal and temporal saccades are included. Note the strong sink in the superficial layers. Right, CSD heatmap of the same animal, but after the silencing. Color scale same as in the center panel. Not the attenuated sink. (**B**) Left, average PETH of discriminating neurons for preferred and non-preferred directions, prior to pulvinar silencing. Right, average PETH of the same neurons, but after pulvinar silencing. *n* = 29, 5 mice. Shaded area, average ± s.e.m. (**C**) Discriminability of the 29 neurons in (B), pre and post silencing of the pulvinar. Gray, individual animals; black, average.

**Fig. S11.**
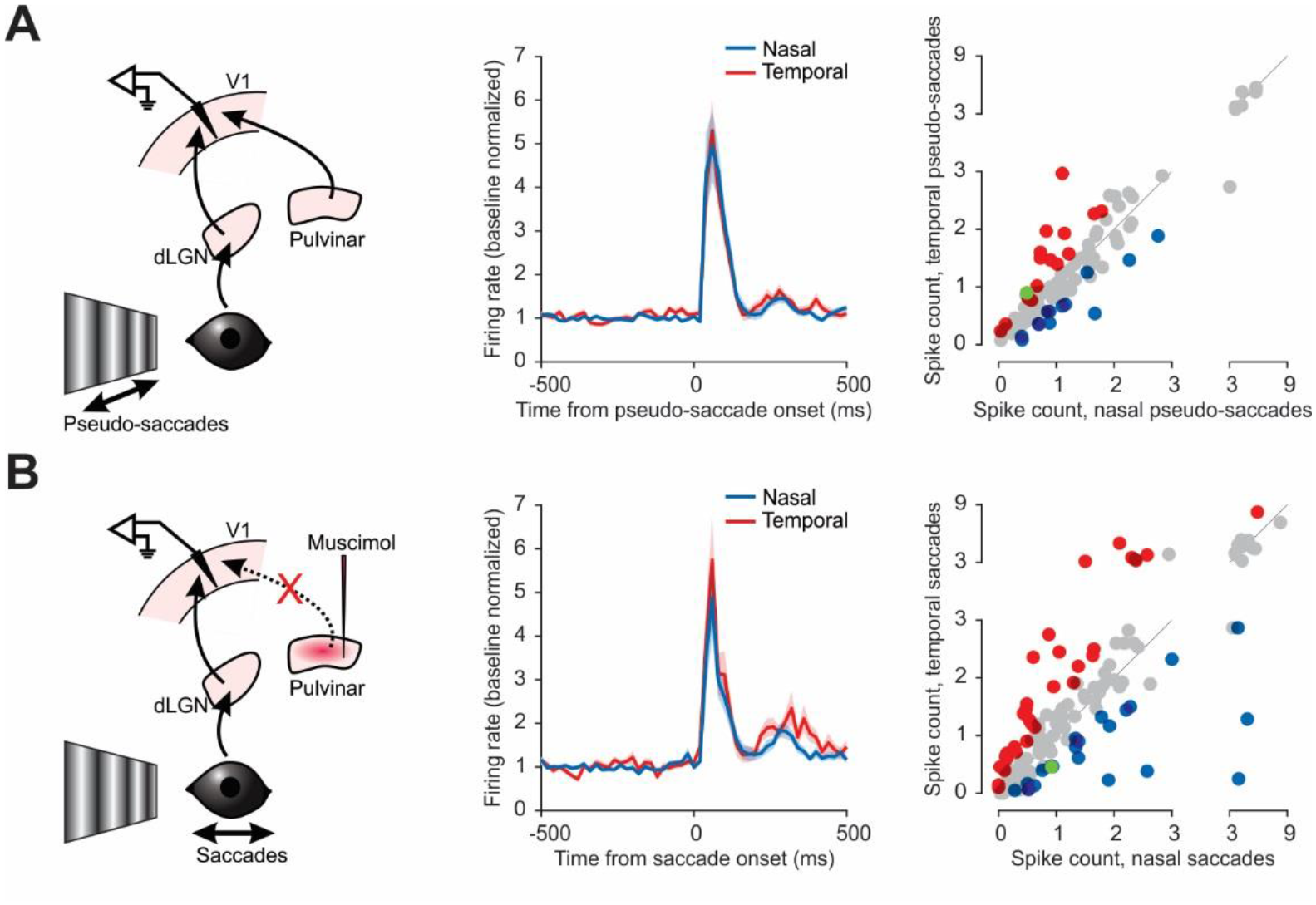
Stimulus direction selective response of V1 neurons in response to saccades under pulvinar silencing and to pseudo-saccades. (**A**) Left, schematic of V1 recording during pseudo-saccades. Center, average PETH for all pseudo-saccade responsive neurons to nasal and temporal directions, baseline normalized (*n* = 102 neurons, 9 mice). Shaded area, average ± s.e.m. Right, scatter plot of the response to nasal and temporal saccades (average spike count in a 100 ms window from saccade onset), for all responsive neurons. Blue, prefer nasal saccades; red, prefer temporal saccades; gray, no statistical difference; green, example neuron in Fig. 5A. (**B**) Same as in (A), but for real saccade responses recorded after pulvinar silencing in the same animals. Center, *n* = 135 neurons. Right, *n* = 53 neurons; green, example neuron in Fig. 5B.

## References

1. T. B. Crapse, M. A. Sommer, Corollary discharge across the animal kingdom. Nat. Rev. Neurosci. 9, 587–600 (2008).

2. M. F. Land, The evolution of gaze shifting eye movements. Curr. Top. Behav. Neurosci. 41, 3–11 (2019).

3. M. F. Land, Eye movements in man and other animals. Vision Res. 162, 1–7 (2019).

4. M. F. Land, Eye movements of vertebrates and their relation to eye form and function. J. Comp. Physiol. A Neuroethol. Sensory, Neural, Behav. Physiol. 201, 195–214 (2014).

5. A. M. Michaiel, E. T. T. Abe, C. M. Niell, Dynamics of gaze control during prey capture in freely moving mice. Elife. 9, 1–27 (2020).

6. A. F. Meyer, J. O’Keefe, J. Poort, Two Distinct Types of Eye-Head Coupling in Freely Moving Mice. Curr. Biol. 30, 2116–2130.e6 (2020).

7. B. Zuber, L. Stark, Saccadic suppression: Elevation of visual threshold associated with saccadic eye movements. Exp. Neurol. 16, 65–79 (1966).

8. M. R. Diamond, J. Ross, M. C. Morrone, Extraretinal control of saccadic suppression. J. Neurosci. 20, 3449–3455 (2000).

9. F. C. Volkmann, Human visual suppression. Vision Res. 26, 1401–1416 (1986).

10. G. W. Beeler, Visual threshold changes resulting from spontaneous saccadic eye movements. Vision Res. 7, 769–775 (1967).

11. E. Matin, Saccadic suppression: A review and an analysis. Psychol. Bull. 81, 899–917 (1974).

12. C. Y. Chen, Z. M. Hafed, A neural locus for spatial-frequency specific saccadic suppression in visual-motor neurons of the primate superior colliculus. J. Neurophysiol. 117, 1657–1673 (2017).

13. A. Thiele, P. Henning, M. Kubischik, K. P. Hoffmann, Neural mechanisms of saccadic suppression. Science. 295, 2460–2462 (2002).

14. F. Bremmer, M. Kubischik, K. P. Hoffmann, B. Krekelberg, Neural dynamics of saccadic suppression. J. Neurosci. 29, 12374–12383 (2009).

15. M. E. Goldberg, R. H. Wurtz, Activity of superior colliculus in behaving monkey. I. Visual receptive fields of single neurons. J. Neurophysiol. 35, 542–559 (1972).

16. F. H. Duffy, C. T. Lombroso, Electrophysiological evidence for visual suppression prior to the onset of a voluntary saccadic eye movement. Nature. 218, 1074–1075 (1968).

17. R. H. Wurtz, Neuronal mechanisms of visual stability. Vision Res. 48, 2070–2089 (2008).

18. D. H. Hubel, T. N. Wiesel, Receptive fields of single neurones in the cat’s striate cortex. J. Physiol. Physiol. 148, 574–591 (1959).

19. D. H. Hubel, T. N. Wiesel, Receptive fields, binocular interaction and functional architecture in the cat’s visual cortex. J. Physiol. 160, 106–154 (1962).

20. C. M. Niell, M. P. Stryker, Highly selective receptive fields in mouse visual cortex. J. Neurosci. 28, 7520–7536 (2008).

21. T. Sakatani, T. Isa, Quantitative analysis of spontaneous saccade-like rapid eye movements in C57BL/6 mice. Neurosci. Res. 58, 324–331 (2007).

22. J. M. Samonds, W. S. Geisler, N. J. Priebe, Natural image and receptive field statistics predict saccade sizes. Nat. Neurosci. 21, 1591–1599 (2018).

23. R. Corazza, C. T. Lombroso, The neuronal dark discharge during eye movements in awake “encéphale isolé” cats. Brain Res. 34, 345–359 (1971).

24. M. Feldman, B. Cohen, Electrical activity in the lateral geniculate body of the alert monkey associated with eye movements. J. Neurophysiol. 31, 455–466 (1968).

25. M. Jeannerod, P. T. S. Putkonen, Lateral geniculate unit activity and eye movements: Saccade-locked changes in dark and in light. Exp. Brain Res. 13, 533–546 (1971).

26. M. M. Roth, J. C. Dahmen, D. R. Muir, F. Imhof, F. J. Martini, S. B. Hofer, Thalamic nuclei convey diverse contextual information to layer 1 of visual cortex. Nat. Neurosci. 19, 299– 307 (2016).

27. H. Nakamura, H. Hioki, T. Furuta, T. Kaneko, Different cortical projections from three subdivisions of the rat lateral posterior thalamic nucleus: A single-neuron tracing study with viral vectors. Eur. J. Neurosci. 41, 1294–1310 (2015).

28. C. Bennett, S. D. Gale, M. E. Garrett, M. L. Newton, E. M. Callaway, G. J. Murphy, S. R. Olsen, Higher-order thalamic circuits channel parallel streams of visual information in mice. Neuron. 102, 477–492.e5 (2019).

29. D. L. Robinson, S. E. Petersen, W. Keys, Saccade-related and visual activities in the pulvinar nuclei of the behaving rhesus monkey. Exp. Brain Res. 62, 625–634 (1986).

30. J. H. Jennings, D. R. Sparta, A. M. Stamatakis, R. L. Ung, K. E. Pleil, T. L. Kash, G. D. Stuber, Distinct extended amygdala circuits for divergent motivational states. Nature. 496, 224–228 (2013).

31. T. K. Sato, M. Häusser, M. Carandini, Distal connectivity causes summation and division across mouse visual cortex. Nat. Neurosci. 17, 30–32 (2014).

32. S.-J. Zhang, J. Ye, C. Miao, A. Tsao, I. Cerniauskas, D. Ledergerber, M.-B. Moser, E. I. Moser, Optogenetic dissection of entorhinal-hippocampal functional connectivity. Science. 340, 1232627 (2013).

33. N. A. Crowder, N. S. C. Price, M. J. Mustari, M. R. Ibbotson, Direction and contrast tuning of macaque MSTd neurons during saccades. J. Neurophysiol. 101, 3100–3107 (2009).

34. D. M. Schneider, A. Nelson, R. Mooney, A synaptic and circuit basis for corollary discharge in the auditory cortex. Nature. 513, 189–194 (2014).

35. J. Yu, D. A. Gutnisky, S. A. Hires, K. Svoboda, Layer 4 fast-spiking interneurons filter thalamocortical signals during active somatosensation. Nat. Neurosci. 19, 1647–1657 (2016).

36. R. Jordan, G. B. Keller, Opposing influence of top-down and bottom-up input on excitatory layer 2/3 neurons in mouse primary visual cortex. Neuron. 108, 1194–1206.e5 (2020).

37. M. Leinweber, D. R. Ward, J. M. Sobczak, A. Attinger, G. B. Keller, A sensorimotor circuit in mouse cortex for visual flow predictions. Neuron. 95, 1420–1432.e5 (2017).

38. W. M. Usrey, S. M. Sherman, Corticofugal circuits: Communication lines from the cortex to the rest of the brain. J. Comp. Neurol. 527, 640–650 (2019).

39. S. M. Sherman, Thalamus plays a central role in ongoing cortical functioning. Nat. Neurosci. 19, 533–541 (2016).

40. R. W. Guillery, S. M. Sherman, Branched thalamic afferents: what are the messages that they relay to the cortex? Brain Res. Rev. 66, 205–19 (2011).

41. R. D. Mooney, S. E. Fish, R. W. Rhoades, Anatomical and functional organization of pathway from superior colliculus to lateral posterior nucleus in hamster. J. Neurophysiol. 51, 407–31 (1984).

42. F. Fredes, T. Vega-Zuniga, H. Karten, J. Mpodozis, Bilateral and ipsilateral ascending tectopulvinar pathways in mammals: A study in the squirrel (spermophilus beecheyi). J. Comp. Neurol. 520, 1800–1818 (2012).

43. N. A. Zhou, P. S. Maire, S. P. Masterson, M. E. Bickford, The mouse pulvinar nucleus: Organization of the tectorecipient zones. Vis. Neurosci. 34, E011 (2017).

44. N. J. Gandhi, H. A. Katnani, Motor functions of the superior colliculus. Annu. Rev. Neurosci. 34, 205–231 (2011).

45. E. Castet, G. S. Masson, Motion perception during saccadic eye movements. Nat. Neurosci. 3, 177–183 (2000).

46. T. L. Watson, B. Krekelberg, The relationship between saccadic suppression and perceptual stability. Curr. Biol. 19, 1040–1043 (2009).

47. T. Watson, B. Krekelberg, An equivalent noise investigation of saccadic suppression. J. Neurosci. 31, 6535–6541 (2011).

48. A. F. Meyer, J. Poort, J. O’Keefe, M. Sahani, J. F. Linden, A Head-Mounted Camera System Integrates Detailed Behavioral Monitoring with Multichannel Electrophysiology in Freely Moving Mice. Neuron. 100, 46–60.e7 (2018).

49. J. S. Stahl, Calcium Channelopathy Mutants and Their Role in Ocular Motor Research. Ann. N. Y. Acad. Sci. 956, 64–74 (2002).

50. B. H. Liu, A. D. Huberman, M. Scanziani, Cortico-fugal output from visual cortex promotes plasticity of innate motor behaviour. Nature. 538, 383–387 (2016).

51. J.. Stahl, A.. van Alphen, C.. De Zeeuw, A comparison of video and magnetic search coil recordings of mouse eye movements. J. Neurosci. Methods. 99, 101–110 (2000).

52. A. Mathis, P. Mamidanna, K. M. Cury, T. Abe, V. N. Murthy, M. W. Mathis, M. Bethge, DeepLabCut: markerless pose estimation of user-defined body parts with deep learning. Nat. Neurosci. 21, 1281–1289 (2018).

53. M. Kleiner, D. H. Brainard, D. G. Pelli, C. Broussard, T. Wolf, D. Niehorster, What’s new in Psychtoolbox-3? Perception. 36, S14 (2007).

54. D. H. Brainard, The Psychophysics Toolbox. Spat. Vis. 10, 433–436 (1997).

55. D. G. Pelli, The VideoToolbox software for visual psychophysics: Transforming numbers into movies. Spat. Vis. 10, 437–442 (1997).

56. G. Bouvier, Y. Senzai, M. Scanziani, Head Movements Control the Activity of Primary Visual Cortex in a Luminance-Dependent Manner. Neuron. 108, 500–511.e5 (2020).

57. M. Pachitariu, N. Steinmetz, S. Kadir, M. Carandini, H. Kenneth D., Kilosort: realtime spike-sorting for extracellular electrophysiology with hundreds of channels. bioRxiv, 061481 (2016).

